# Cryo-EM structure of the complete and ligand-saturated insulin receptor ectodomain

**DOI:** 10.1101/679233

**Authors:** Theresia Gutmann, Ingmar Schäfer, Chetan Poojari, Beate Brankatschk, Ilpo Vattulainen, Mike Strauss, Ünal Coskun

**Affiliations:** Paul Langerhans Institute Dresden of the Helmholtz Zentrum Munich at the University Hospital and Faculty of Medicine Carl Gustav Carus of TU Dresden, Dresden, Germany; German Center for Diabetes Research (DZD e.V.), Neuherberg, Germany; Department of Structural Cell Biology, Max Planck Institute of Biochemistry, Munich, Germany; Department of Physics, University of Helsinki, P.O. Box 64, FI-00014, Helsinki, Finland; Computational Physics Laboratory, Tampere University, Tampere, Finland; Department of Anatomy & Cell Biology, McGill University, Montreal, Quebec, Canada

## Abstract

Glucose homeostasis and growth essentially depend on the peptide hormone insulin engaging its receptor. Despite biochemical and structural advances, a fundamental contradiction has persisted in the current understanding of insulin ligand–receptor interactions. While biochemistry predicts two distinct insulin binding sites, 1 and 2, recent structural analyses have only resolved site 1. Using a combined approach of cryo-EM and atomistic molecular dynamics simulation, we determined the structure of the entire dimeric insulin receptor ectodomain saturated with four insulin molecules. Complementing the previously described insulin–site 1 interaction, we present the first view of insulin bound to the discrete insulin receptor site 2. Insulin binding stabilizes the receptor ectodomain in a T-shaped conformation wherein the membrane-proximal domains converge and contact each other. These findings expand the current models of insulin binding to its receptor and of its regulation. In summary, we provide the structural basis enabling a comprehensive description of ligand–receptor interactions that ultimately will inform new approaches to structure-based drug design.

**In brief:** A cryo-EM structure of the complete insulin receptor ectodomain saturated with four insulin ligands is reported. The structural model of the insulin–insulin receptor complex adopts a T-shaped conformation, reveals two additional insulin-binding sites potentially involved in the initial interaction of insulin with its receptor, and resolves the membrane proximal region.

## Introduction

The insulin receptor (IR) signaling system is a key regulator of metabolism and cellular growth. Its dysfunction is linked to clinical manifestations such as diabetes mellitus, cancer, and Alzheimer’s disease (Saltiel and Kahn, 2001; Belfiore and Malaguarnera, 2011; Kleinridders et al., 2014). The IR is an extensively glycosylated disulfide-linked (αβ)_2_ homodimer with a modular domain structure (Fig. 1A). Each protomer consists of an extracellular ligand-binding α subunit and the membrane-spanning β subunit, which also harbors the intracellular kinase domain. The modular organization of the ectodomain (ECD) with high intrinsic flexibility poses a challenge to structural studies of the IR, as do the branched sugars of the glycosylation sites, and its complex ligand binding properties. Insulin binding to the ectodomain concomitantly elevates the receptor’s intrinsic tyrosine kinase activity prior to cellular signal transduction (Kasuga et al., 1982). The precise mechanism of how insulin initially engages its receptor, as well as the associated conformational changes leading to tyrosine kinase signaling still remain elusive (De Meyts, 2015; Tatulian, 2015).

**Fig. 1:**
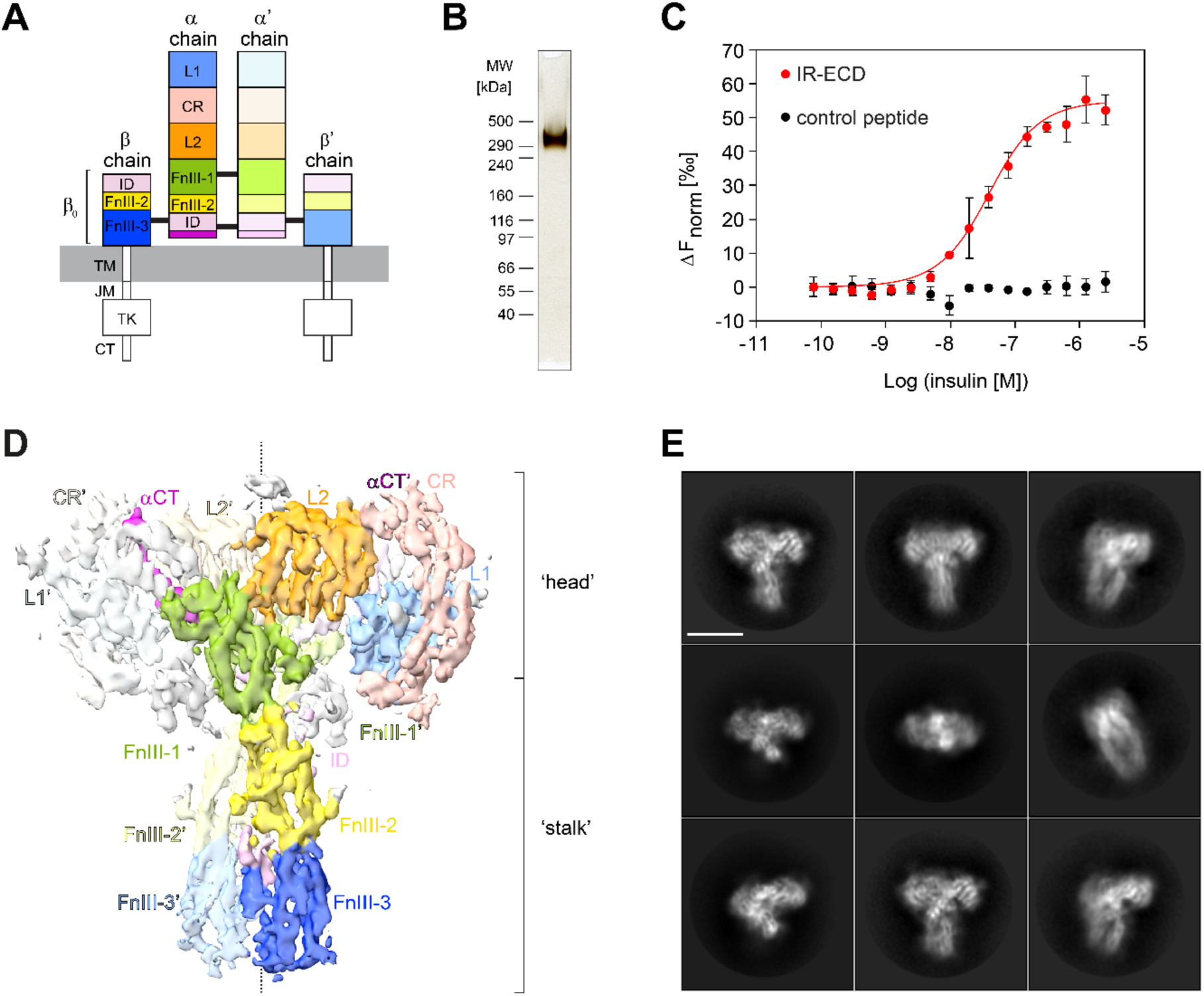
Insulin receptor ectodomain purification and cryo-EM. (**A**) Scheme of insulin receptor domain architecture: L1 and L2, leucine-rich repeat domains 1 and 2; CR, cysteine-rich domain; FnIII-1,-2,-3, fibronectin type-III domains 1, 2, 3; ID, insert domain; TM, transmembrane; JM, juxtamembrane; TK, tyrosine kinase domain; CT, C-terminal tail. The alpha C-terminal regions (αCT, αCT’) are drawn in purple. Black lines indicate inter-subunit disulfide bonds. A prime (’) denotes the chain, domain, or residue within the second protomer. (**B**) Purified dimeric IR-ECD (IR(αβ_0_)_2_) migrates as a single band with an apparent molecular weight of 351 kDa on a non-reducing 3-8% Tris-Acetate SDS-PAGE gel as visualized by silver staining. (**C**) Equilibrium binding to native human insulin in solution was assessed by microscale thermophoresis of IR-ECD after tris-NTA-RED labelling. An 8xHis-tagged control peptide served as negative control to rule out unspecific binding or interference with the tris-NTA-RED dye. The normalized fluorescence difference (ΔF_norm_) is plotted against ligand concentration. Error bars display standard deviations; n = 3. (**D**) Front view of the IR-ECD cryo-EM density map saturated with insulin ligands at 4.3 Å estimated nominal global resolution. Domains are colored as in (A). (**E**) Representative 2D class averages of particles contributing to the reconstruction in (D) of the IR-ECD exposed to human insulin. The scale bar corresponds to 10 nm.

Crystallography of the unliganded (i.e. *apo*) IR-ECD dimer has revealed a structure resembling an inverted ‘U’ or ‘V’ with respect to the membrane, placing the membrane insertion sites ∼ 11.5 Å apart from each other (McKern et al., 2006; Croll et al., 2016). Single-particle EM of full-length IR in lipid nanodiscs corroborated that this *apo*-conformation is retained in the membrane context. Insulin binding converts the receptor ectodomain into a T-like shape that draws the membrane-proximal fibronectin domains closer together, enabling transmembrane signaling (Gutmann et al., 2018). Due to the low resolution of the negative-stain 2D class averages, no structural information about the location and number of bound insulin molecules could be obtained. The T-shaped conformation was confirmed shortly after by cryo-EM of the IR-ECD in complex with one or two insulins bound to the N-terminal domains (Scapin et al., 2018). However, major parts of the fibronectin domains could not be reconstructed, preventing conclusions on the transmembrane signalling mechanism.

In another cryo-EM approach, the soluble ectodomain was fused to a C-terminal leucine zipper in an attempt to reduce conformational heterogeneity and mimic membrane anchorage, thus restoring insulin-binding properties of the complete receptor (Hoyne et al., 2000; Weis et al., 2018). Structural heterogeneity was further decreased by mild deglycosylation and complexation with Fv variable domain modules of the anti-IR antibody 83-7. These modifications enabled the capture of a singly liganded transition state with insulin bound to the N-terminal region and the fibronectin regions in a pincer-like fashion (Weis et al., 2018).

Previous biochemical and mutagenesis experiments have mapped two distinct binding sites, termed sites 1 and 2, on both the IR and on insulin (De Meyts et al., 1978; De Meyts, 2015). While site 1 ligand receptor interactions were largely confirmed (Menting et al., 2013; Menting et al., 2014; Scapin et al., 2018; Weis et al., 2018), site 2 interactions could not be demonstrated structurally.

Here, by applying single-particle EM and atomistic molecular dynamics (MD) simulations, we report the structure of the complete, pseudo-symmetric human insulin receptor ectodomain in a T-like conformation saturated by four insulins. We observe that the membrane-proximal fibronectin domains converge, highlighting the coupling of ligand binding and fibronectin domain interactions as intrinsic features of the IR-ECD. While two of the observed insulin binding sites agree with those mapped in the ‘head’ region (Scapin et al., 2018; Weis et al., 2018), the additional two insulin molecules are located in the now fully-resolved ‘stalk’ regions (Fig. 1D), providing unambiguous structural evidence for the existence and mechanism of site 2 binding.

## Results and discussion

### Purification and biochemical characterization of the complete human IR ectodomain

The complete insulin receptor ectodomain (IRαβ_0_) (Fig. 1A) was produced by secretion from human embryonic kidney cell-derived cells, ensuring human-like post-translational processing, such as glycosylation. Purification of the recombinant protein directly from the medium resulted in a highly pure and ligand-sensitive receptor ectodomain that was amenable to cryo-EM studies. SDS-PAGE confirmed a complete, glycosylated, dimeric polypeptide composition with an apparent molecular weight of 351 kDa (Fig. 1B, S1). Ligand binding was assessed by two independent assays in solution (Fig. 1C, S2). In both assays, ligand labeling was entirely omitted in order to preserve binding properties. First, the thermal stability of purified IR-ECD was followed by low volume differential intrinsic tryptophan scanning fluorimetry (nano-DSF) with a temperature gradient from 20 °C to 95 °C (Fig. S2A). In the absence of insulin, IR-ECD unfolds in two steps with transition temperatures of 58.9 °C and 64.0 °C. Interestingly, insulin binding shifted only the first transition temperature down to 51.1°C, implying that insulin binding leads to conformational changes within distinct regions of the IR-ECD. Next, ligand binding affinity assessed by microscale thermophoresis (Seidel et al., 2013) showed an EC_50_ = 39.5 ± 9.4 nM (Fig. 1C, S2C), corresponding to a low-affinity binding regime. This is in good agreement with the established concept that without membrane anchorage the IR-ECD loses high-affinity binding in the subnanomolar range (∼ 0.2 nM) (Whittaker et al., 1994; Bass et al., 1996; Whittaker et al., 2008; Subramanian et al., 2013; De Meyts, 2015), similar to the EGF receptor ectodomain (Lax et al., 1991; Ferguson et al., 2003).

### Single-particle cryo-EM analysis of the IR-ECD

The IR-ECD was analyzed by single-particle cryo-EM in the absence and presence of recombinant human insulin. Vitrification conditions allowed cryo-EM data collection for the unliganded and liganded ectodomain. In the absence of ligand, 2D class averaging revealed considerable structural heterogeneity (Fig. S3). Although individual domains could be identified in a subset of 2D class averages, no high-resolution features, such as clearly identifiable, individual secondary structural elements, were apparent. Consequently, attempts at reconstructing these data in 3D did not yield any sub-nanometer EM maps. In particular, the fuzzy haphazard appearance of membrane-proximal fibronectin domains in the 2D class averages points at considerable flexibility. This behavior of the IR-ECD in isolation most likely reflects the presence of various transition states and conformations sampled in the absence of insulin.

For cryo-EM samples of the liganded IR-ECD, saturating amounts of 50 μM insulin were used, corresponding to a receptor:ligand molar ratio of 1:35. The rationale for such a large ligand excess was to ensure saturation of all available insulin binding sites on the receptor and the associated reduction of structural heterogeneity. Now, individual secondary structure elements became clearly discernable after 2D classification (Fig. 1, S4, S5). After further classification steps, 3D refinement, B-factor weighting, and map sharpening, the 3D reconstruction of the liganded IR-ECD reached an apparent overall resolution of 4.3 Å, as estimated by the Fourier shell correlation of independently refined half-maps (0.143 criterion (Rosenthal and Henderson, 2003)) (Fig. S4, S5). Our 3D reconstruction confirmed the T-like structure as seen in the insulin-bound full-length IR at low resolution by negative stain EM (Gutmann et al., 2018). Features of the compact head containing L1, CR, and L2 domains appear better defined than the fibronectin stalks, which exhibit more flexibility (cp. local resolution estimate in Fig. S5F). Importantly, we have refrained from applying C2 symmetry during any of the processing steps, in line with the observed inherent asymmetry in the liganded IR-ECD dimer (Fig. S5). This asymmetry, as with the flexibility, is most pronounced in the membrane-proximal stalks (Fig. S6).

All structured domains of the IR-ECD, as well as the localization of the insulins, were unambiguously identified in our cryo-EM density map (Fig. 1D, 2 and Tab. S1, S2). This map in combination with previously published structural information enabled us to construct a single model for the nearly complete ECD in complex with four insulins (Fig. 2). The single exceptions were the intrinsically disordered insert domains (ID) of the IR-ECD that could only partially be modeled into certain incohesive density features in the vicinity of the fibronectin domains. The IDs encompass the cleavage site for furin (720-723 Arg-Lys-Arg-Arg), which processes the IR(αβ) polypeptide chain into the IRα and IRβ chains. Thus, the ID is separated into IDα and IDβ in the mature receptor dimer. Our data allowed us to tentatively model the IDα loop but we refrained from including IDβ in the cryo-EM structure. To err on the side of caution, we did not model the furin cleavage sites in the cryo-EM structure, even though a non-contiguous density feature attributable to this part of the IR-ECD α’-chain is present in the map (Tab. S2).

**Fig. 2:**
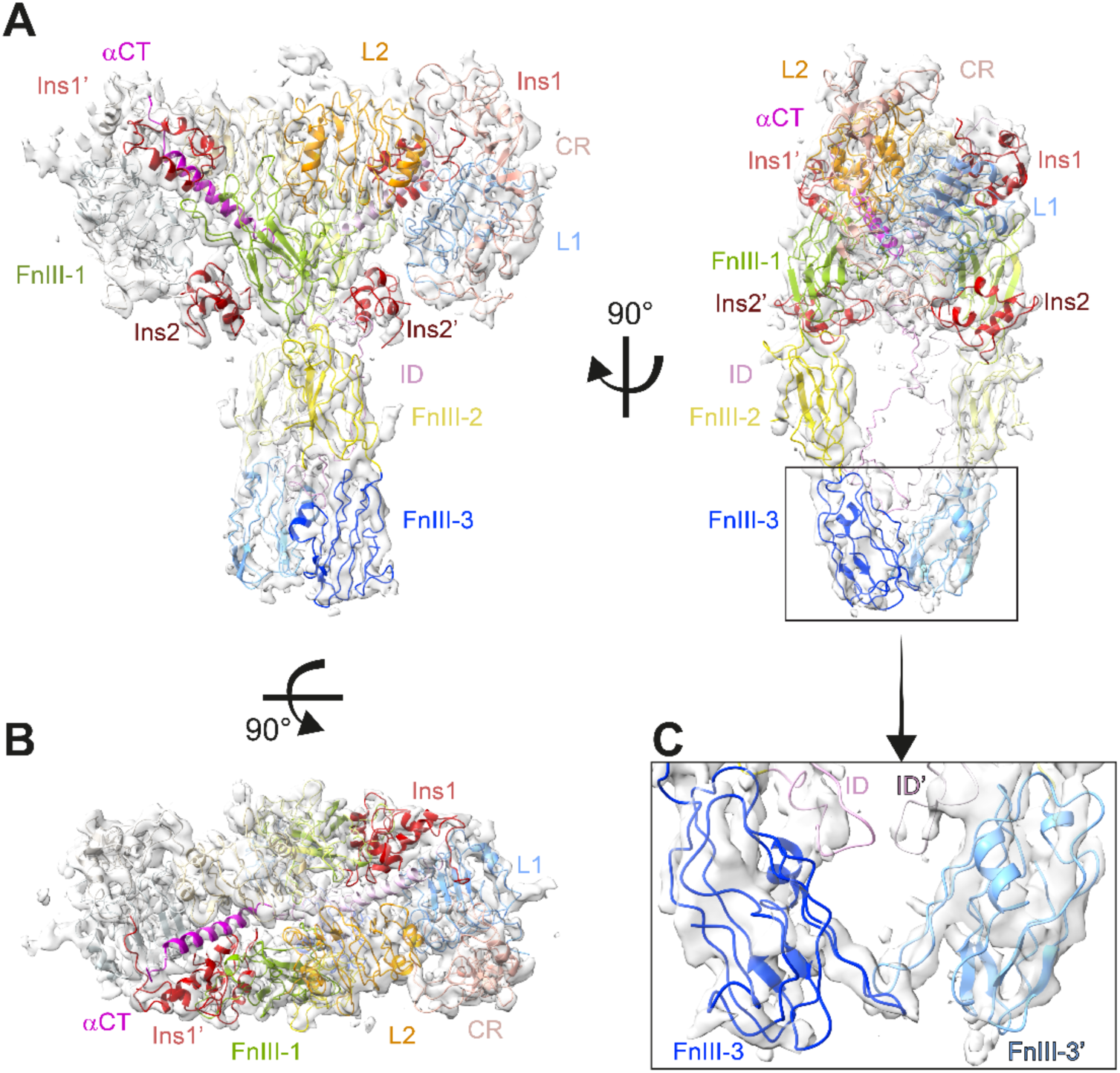
Cryo-EM structure of the ligand-saturated insulin receptor ectodomain. (**A**, **B**) Orthogonal views of the cryo-EM map and structure of the IR-ECD dimer complex. (**C**) Close-up of the membrane-proximal FnIII-3 domains. The color code for the individual domains in all panels is as in Fig. 1; the four insulin moieties are depicted in red.

After initial rigid body docking and local flexible fitting of the IR-ECD domains into the EM map, the resulting structure was manually rebuilt and refined. The final model conforms to commonly accepted quality indicators (Tab. S1). For atomistic MD simulations, we completed our cryo-EM structure by appending all unresolved loops (Tab. S2) and glycosylations (Sparrow et al., 2007; Sparrow et al., 2008) (Fig. S7, Tab. S3).

### Conformation of the insulin-saturated IR-ECD

In accord with previous data, significant conformational changes in the insulin-bound IR-ECD are observed with respect to rigid-body rotations in the *apo*-IR-ECD (McKern et al., 2006; Croll et al., 2016; Gutmann et al., 2018; Scapin et al., 2018). The liganded IR-ECD adopts a T-shaped conformation similar to the membrane-embedded full-length receptor at low resolution (Fig. 1D, 2) (Gutmann et al., 2018). As mentioned above, our model was extended to include the stalks, resolving the entire IR-ECD. The fibronectin domains come together in a pincer-like fashion (Fig. 2C) similar to what has been described for the C-terminally tethered IRΔβ-zipInsFv (Weis et al., 2018). Although the arrangement of fibronectin domains differs from that in IRΔβ-zipInsFv, proximity of the same FnIII-3 domain loops was reproduced in our cryo-EM structure and MD simulations (in particular residues Asp854-His858, Fig. S8). Thus, the membrane-proximal domains are capable of interacting in the absence of a C-terminal zipper element. The interaction between the FnIII-3 and FnIII-3’ domains further supports the concept that receptor activation is directly linked to the lateral distance between transmembrane domains (and consequently the relative orientation of the attached intracellular kinase domains) (Kavran et al., 2014; Gutmann et al., 2018).

### Structural characterization of insulin binding to sites 2 and 2’

Earlier biochemical studies showed that each insulin receptor protomer contains two distinct insulin-binding sites, termed site 1 and 2 (site 1’ and 2’ on the other protomer) (De Meyts, 1994; Schaffer, 1994; De Meyts, 2015). The IR-ECD head region of our structural model agrees well with the IR-ECD cryo-EM structure in complex with two insulins in site 1/1’ (Scapin et al., 2018): the L1-CR-L2 module is complexed by one insulin and adopts a ∼ 90° angle with respect to the [L2-(FnIII-1):L2’-(FnIII-1’)] module, and insulin 1 interacts with the L1-αCT’ tandem element and loops of the FnIII-1’ domain (L1’+αCT and FnIII-1 in case of insulin 1’) (Fig. 2). Contrasting to previous observations, two additional, previously undescribed density features contacting the FnIII-1 and FnIII-1’ domains were identified in our cryo-EM map (Fig. 1D, 2). These features did not correspond to unmodeled regions of the IR-ECD but could be clearly assigned as additional insulin molecules (Fig. 2, 3). The concordance between these density features and insulin is further supported by the observation that the map-versus-model correlation coefficient of both head- and stalk-bound insulins is almost indistinguishable. If anything, it is slightly inferior for the head-bound ligands (Tab. S1). These two additional insulin moieties, analogously named insulin 2 and 2’, interact with β-sheets and inter-strand loops (Tyr477-Trp489, Asp535-Arg554) of the FnIII-1 and FnIII-1’ domain, respectively (Fig. 2, 3, S9). Among those residues, Lys484 and Leu552 have been implicated in insulin binding to site 2 in the holoreceptor by alanine scanning mutagenesis screens (Whittaker et al., 2008). Also, insulins 2 and 2’ interact with a loop within the L1’ (Asp151’-Glu154’) and L1 domain (Asp151-Glu154), respectively (Fig. S9).

**Fig. 3:**
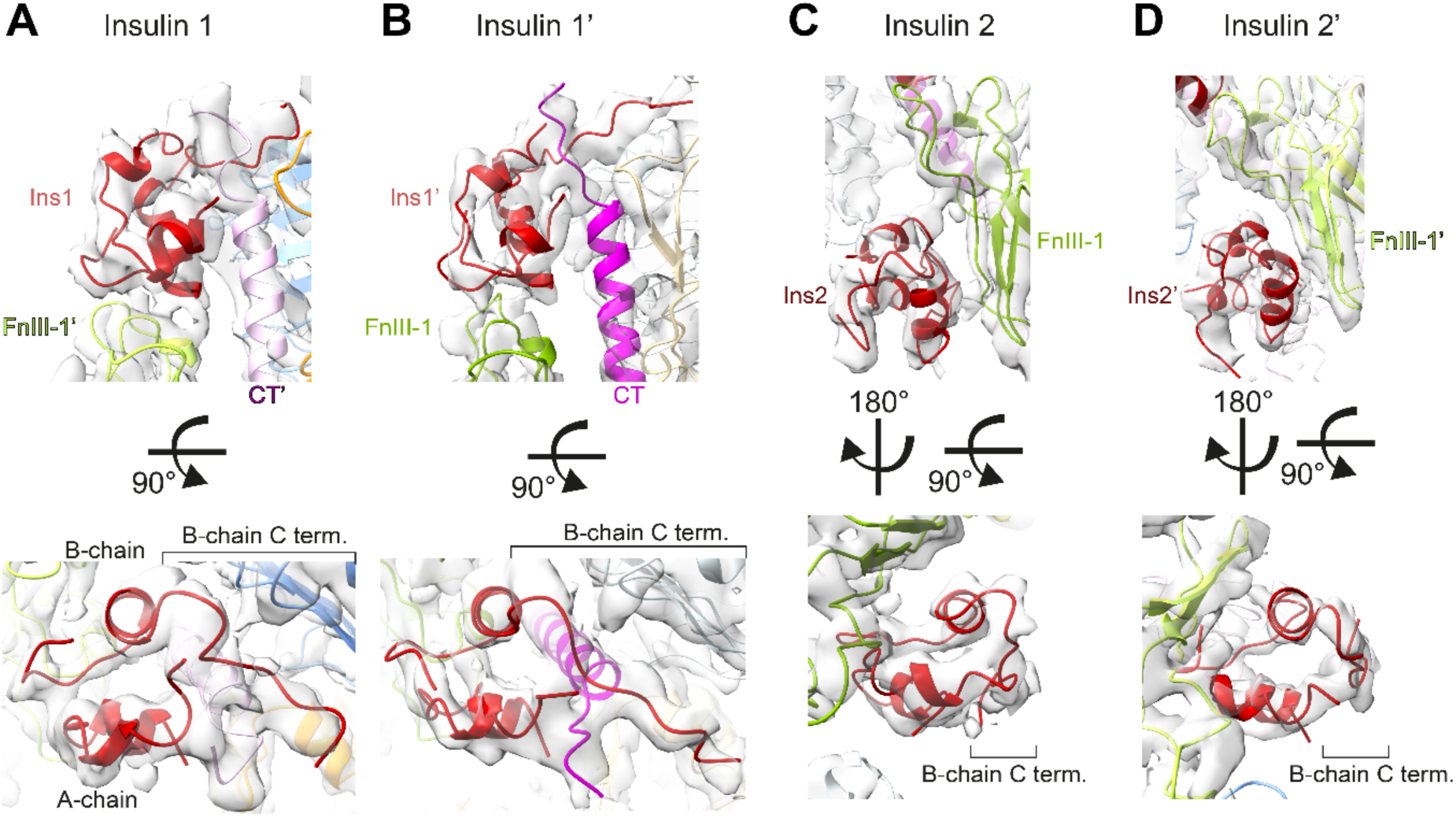
Binding sites and conformations of insulins bound to IR-ECD. The cryo-EM density map and structure in close-up views of the four insulins and their receptor binding sites are displayed: (**A**) insulin 1, (**B**) insulin 1’, (**C**) insulin 2, and (**D**) insulin 2’. Insulin structures are aligned below for comparison illustrating the open conformation of site-1-bound insulins with a detached C-terminal B-chain segment in contrast to the closed conformation at site 2. Individual domains in all panels are colored as in Fig. 1 and 2.

Interestingly, head- and stalk-bound insulins display different conformations. Insulin consists of two polypeptide chains, an A-chain of 21 residues structured into two α-helices separated by a stretch of extended polypeptide, and a B-chain of 30 residues with a central α-helix (De Meyts, 2015). Insulins 1 and 1’ were in the receptor-bound open conformation with a detached B-chain C terminus (including the aromatic triplet Phe B24, Phe B25, and Tyr B26) that is critical for engaging receptor site 1 (Hua et al., 1991; Menting et al., 2014). In contrast, insulins 2 and 2’ have a ‘closed conformation’, which corresponds to the conformation in solution prior to IR binding (Hua et al., 1991; Menting et al., 2014) (Fig. 3, bottom panels).

### Atomistic MD simulations of insulin-saturated IR-ECD

To follow the dynamics of insulin:IR-ECD interactions with high spatial and temporal resolution, we performed atomistic MD simulations. For the sake of completeness, we extended our experimentally determined insulin–IR-ECD model by incorporating the previously absent IDβ loops, as well as the N- and O-linked glycans (Fig. S7, Tab. S3). Since furin cleaves C-terminally of this sequence and because there is no evidence of its removal in the secreted ectodomain, we also included the furin cleavage site (Arg-Arg-Lys-Arg) to the αCT helix in our simulation models. In fact, in the case of αCT’ the furin cleavage site could be fitted into our density map; however, the resolution in this region was too low to finalize this part of the model (Tab. S2).

In 10 independent 500 ns simulations, the fully liganded ECD displayed a rather rigid head region and more flexible stalks (Fig. S6B), in line with our cryo-EM data. In all simulations, the four insulins remained bound to the receptor. Here, a contact between two residues was considered to be established as a stable interaction if the distance between any pair of atoms in the two residues was ≤ 3.5 or ≤ 6 Å, and the occupancy at this distance was at least 50% (Fig. S10, S11). Three key areas of interaction stabilizing insulin 2 (or 2’) could be discerned. First, interactions formed between the central A-chain insulin residues (Ile A10-Glu A17) and FnIII-1 domain residues (Leu486-Arg488, Asp535-Leu538, Asn547-Leu552), second, the interactions between N-terminal residues of the insulin B-chain (Phe B1-Leu B6) and FnIII-1 domain residues (Trp551-Arg554) and third, those between B-chain α-helix residues (His B10-Glu B21) and FnIII-1 domain residues (Tyr477-Arg488, Leu552-Arg554). Interestingly, insulins 2 and 2’ display some asymmetry in their binding. In particular, insulin 2’ appears to additionally interact with both ID loops (Gln 672-Ser673 and Glu676’-Cys682’). Insulin 2’-ID interactions were preserved across all 10 MD simulations. Moreover, insulin 2 appears to interact with its residues (Gly A1, Glu A4, Gln A5, Thr A8, Ser A9) with a single L1’ domain loop (Asp151’-Glu154’). In case of insulin 2’ only Gln A5 is in contact with L1 (Asn152 and Glu154). Stalk insulins 2/2’ are localized mainly pack against β-strands of FnIII-1/1’ domains, which provide a stable interface for binding. In the case of head-bound insulins, the interaction site is composed of flexible loops from several domains. Thus, the binding of insulin 1 and 1’ depends on the proper organization of all required domains and is hence more sensitive to conformational variations.

Fluctuations within the insulin conformations during our simulations are mostly attributed to the flexible A- and B-chain termini as indicated in the root-mean-square deviation (RMSD) plots (Fig. S12, S13). In particular, the insulin B-chain non-helical N and C termini featured notable flexibility in our simulations (Fig. S13). C-terminal B-chain dynamics were measured here based on center-of-mass distances between the Cα atoms of the C-terminal residues (Phe B24-Thr B30) and B-chain α-helix residues (Gly B8, Val B12, and Leu B15) (Fig. S14, Tab. S4). The C-terminal B-chain fluctuations were most pronounced in insulins 2 and 2’, which, unlike insulins 1 and 1’, do not engage their B-chain C termini with the receptor.

## Discussion

Our cryo-EM structure of the human IR-ECD in complex with four insulin molecules offers new insights into the full IR-ECD in its ligand-saturated state and helps to reconcile a number of earlier findings to inform an integrated model of insulin–IR binding and activation.

Insulin binding to the full-length receptor is characterized by high and low-affinity binding and/or negative cooperativity (De Meyts, 1994; Whittaker et al., 2008). Based on photo-crosslinking and mutagenesis screens, two distinct molecular surfaces on the insulin molecule have been identified to interact with two distinct receptor sites (1/1’ and 2/2’) (De Meyts, 1994; De Meyts, 2015). It was furthermore proposed that insulin cross-links both receptor monomers by binding in a bivalent manner (De Meyts, 1994; De Meyts, 2015). The interactions of insulin with site 1 (and 1’) on the IR-ECD agree very well with previously described structures of insulin bound to the classical binding site 1 / 1’ (Menting et al., 2013; Menting et al., 2014; Scapin et al., 2018; Weis et al., 2018). The classical insulin site 1 residues, which are widely conserved during vertebrate evolution participated in this interaction in our structure and in MD simulations (i.e. Gly A1-Glu A4, Tyr A19, Asn A21, Gly B8, Ser B9, Leu B11, Val B12, Tyr B16, Phe B24, Phe B25, Tyr B26) (Fig. 4, S9–11). Insulin bound to site 1 (or 1’) simultaneously interacts with residues from the FnIII-1’ (or FnIII-1) domains, similar to the very recent IR-ECD cryo-EM structures (Scapin et al., 2018; Weis et al., 2018), supporting the earlier proposed bivalent insulin binding mode. The structural basis for site 2 insulin interactions, however, remained elusive and an irreconcilable difference between biochemical studies and previous cryo-EM data available for insulin–IR engagement persisted.

**Fig. 4:**
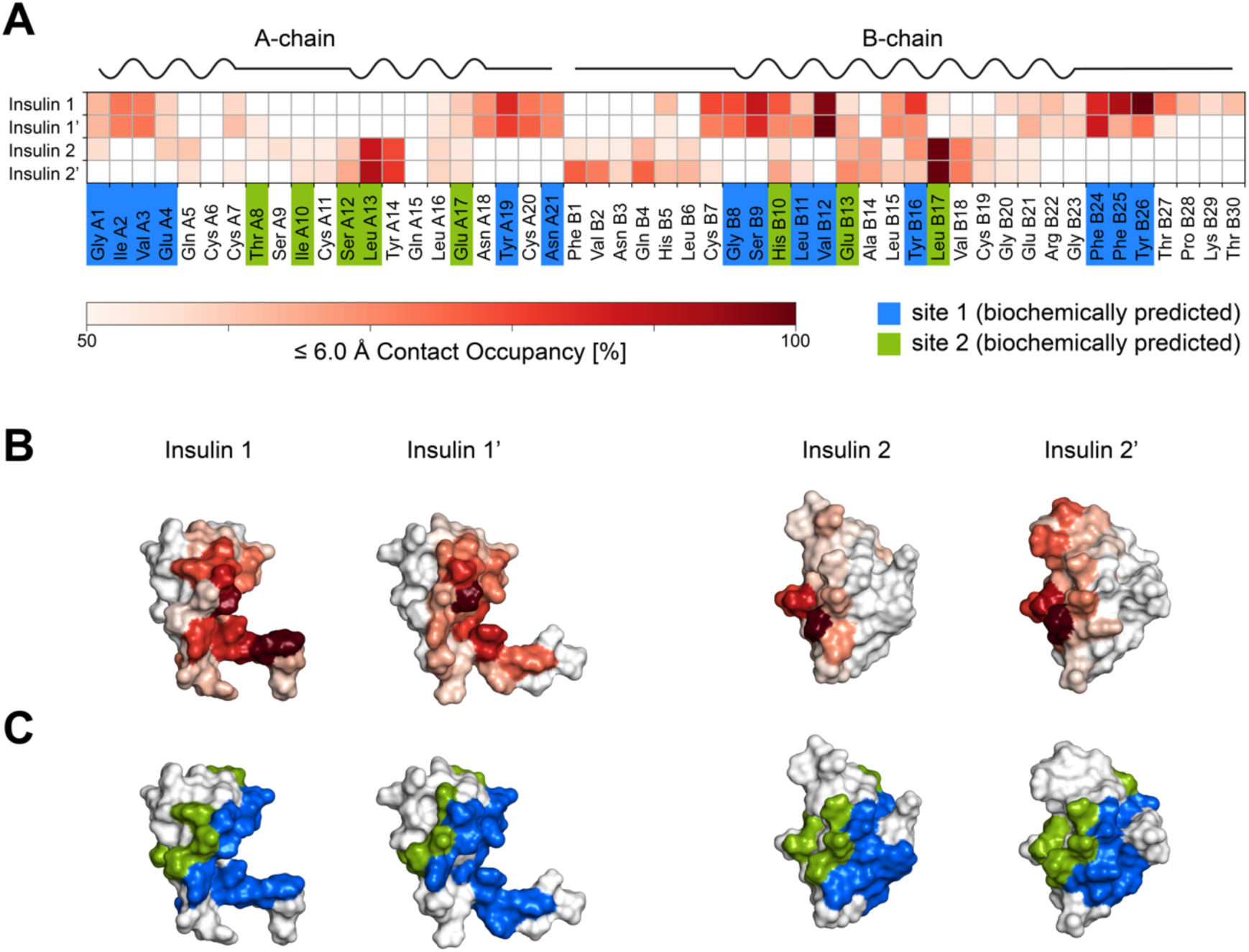
Interactions of insulin with the IR-ECD binding sites 1 and 2. (**A**) Summary of per residue contact occupancies of the four insulins bound to IR-ECD in the MD simulations. The contact occupancies are encoded by shades of red. Only contacts of at least 50% occupancy are displayed. Residues shown previously to contribute to interactions with IR-ECD site 1 and site 2 in biochemical experiments are highlighted in blue and green, respectively. The secondary structure of human insulin is drawn schematically above the contact map (based on PDB:3I3Z). (**B**) Contact occupancies color-coded as in (A) are plotted on the structural models of the four insulins bound to the IR-ECD. This is juxtaposed with coloring of biochemically predicted site 1 and site 2 residues in blue and green, respectively, in (**C**).

The biochemically mapped sites 2 and 2’ involve residues in the IR-ECD FnIII-1 domains (De Meyts et al., 1978; Whittaker et al., 2008; De Meyts, 2015; Ye et al., 2017), which agree well with our structural data. All insulin residues proposed to be involved in site 2 contacts participated in this interaction, i.e. Thr A8, Ile A10 Ser A12, Leu A13, Glu A17, His B10, Glu B13, and Leu B17 (Fig. 4).

### What is the role of the interaction of insulin with sites 2 and 2’?

While the stalk-bound insulin molecules in our structure display a ‘closed’ conformation reminiscent of insulin conformation in solution prior to binding (Hua et al., 1991), the head-bound insulins adopt an ‘open’ conformation, previously described upon receptor site 1 binding (Xu et al., 2009; Menting et al., 2014; Weiss and Lawrence, 2018). This suggests that site 2 (or 2’) interactions might be important for establishing the initial ligand–receptor contact and possibly contribute to ligand specificity. The arrangement of sites 1 and 2’ in the *apo*-IR-ECD crystal structure also implicates sites 2 and 2’ as the sites of first contact. In this structure, sites 2 and 2’ both are exposed and solvent-accessible (McKern et al., 2006; Whittaker et al., 2008; Croll et al., 2016), while residues in sites 1 and 1’ appear partially engaged in interactions with the opposite FnIII-2 domain (Fig. S15). Earlier biochemical findings had already indicated site 2 as the initial insulin contact site (De Meyts, 2004). Evolutionarily ancient vertebrate insulin from hagfish (*Myxine glutinosa*) was shown to exhibit anomalous binding behavior, different from most mammalian insulins. Despite absolute conservation of all site 1 residues and structural homology, hagfish insulin displays slow association kinetics, low affinity, low metabolic potency, and decreased negative cooperativity (Muggeo et al., 1979; De Meyts, 2004). This was attributed to variations in site 2 residues Leu A13 and Leu B17, which contribute significantly to site 2 interactions in our data (Fig. 4). Analogously, insulin variants carrying alanine replacements of those residues display a 20-fold decrease in initial ligand–receptor association (DeMeyts et al., 1976; De Meyts and Whittaker, 2002).

Based on the above-described structural and biochemical findings by us and others, a molecular model is envisaged where insulin initially interacts with one of the exposed sites 2. This may destabilize certain *apo*-conformation-specific, inhibitory domain-domain interactions such as the L1:FnIII-2’ cross-talk, which in turn permits T-shape specific domain-domain interactions. The resulting transition states may be similar to those captured in the cryo-EM data for unliganded IR-ECD (Fig. S3). The initial capture of insulin by site 2 may also relate to a conformational change important for ligand engagement with site 1. Indeed, the C-terminal segments of the insulin B-chains in both insulins 2 and 2’ were not involved in site 2 interactions at any time in our MD simulations, but seemed to sample their environment comparatively unrestrained. In a situation where site 2’ is liganded and site 1 is still in a transition state, these C termini might be free to establish contacts with site 1 adopting an ‘open conformation’. The possibility that these segments play a critical role in initiating a series of conformational changes was suggested very recently (Weiss and Lawrence, 2018). Finally, the receptor binding sites 1 and 1’ on the head are fully formed and insulin interacts with both receptor monomers in a bivalent manner (De Meyts, 1994; De Meyts, 2015).

In the presence of micromolar insulin concentrations, negative cooperativity is reversed and remaining free binding sites are believed to become saturated, inducing the so-called slowly dissociating (*K*_super_) state (De Meyts and Whittaker, 2002; Kiselyov et al., 2009). Under these conditions, site 2-bound insulins may assume yet another function by acting as a molecular wedge to prevent L1 from folding back onto FnIII-1/2’, thus stabilizing the T-shaped conformation. This view is also supported by our observation of L1/L1’ interacting with insulin 2’/2. Strikingly, the stalk-binding sites are partially asymmetric in our MD simulations: only one insulin molecule (insulin 2’ bound to FnIII-1’) appears to interact with residues of the ID regions of both monomers. It is therefore tempting to speculate whether this asymmetry-inducing interaction is critical for generating negative cooperativity in the cell surface receptor. Such an interaction would likely influence the positioning of the αCT/ αCT’ helices critical for high-affinity binding and for cross-talk with the stalks. Another potentially cross-talking element might be the FnIII-1 domains, which feature one of the inter-protomer disulfide bonds and contribute to binding of insulin to sites 1 / 1’ and 2 / 2’ (Fig. 3).

It is not known whether initial insulin docking at the receptor induces a conformational change or whether the receptor transiently adopts various conformations allowing the ligand to engage, or both. The high structural heterogeneity of the unliganded ectodomain observed here (Fig. S3) supports the view that the insulin-free IR adopts various transition states and thereby samples its environment. Even though this interpretation is in line with earlier *in silico* predictions, the precise contribution of distinct receptor conformations to signaling remains to be understood in detail (Kiselyov et al., 2009). This is not to imply that the four-insulin-bound state reported here necessarily corresponds to the most prevalent, physiologically active IR conformation. However, this structure exhibits all possible binding sites including those important for initial contact, albeit with possibly differing precise binding modes in the transition states.

Although insulin replacement remains an essential therapy, it is still hampered by the inability of exogenously administered insulins to recapitulate the full spectrum of physiological insulin action (Jiracek and Zakova, 2017). A thorough understanding of the molecular details of ligand–insulin receptor activation is prerequisite for the development of specific agonists as well as antagonists. We and others have suggested that the transmembrane signaling mechanism of the IR relies on the control of the distance between the transmembrane domains exerted by this ectodomain (Whittaker et al., 1994; Kavran et al., 2014; Gutmann et al., 2018). Similar to the IR, the distance between the membrane-proximal regions of the structurally related mitogenic IGF1 receptor ectodomain is reduced from ∼ 115 to ∼ 67 Å in its unliganded form (Xu et al., 2018). This implies that the fine tuning of transmembrane domain positioning and orientation may be an intricate detail of ligand selectivity and cell signaling outcomes via allosteric domain coupling across the membrane. Therefore, integrating membrane lipid composition and lipid–protein interactions at the next level of reconstruction and analysis will indispensably contribute to a complete understanding of receptor function and improved pharmacological targeting (Coskun and Simons, 2011; Endres et al., 2014; Kaszuba et al., 2015).

## Materials and methods

### Cloning and production of IR-ECD

A gene encoding IR-ECD (IR signal sequence followed by residues 1-917 of the mature IR isoform A (uniprot entry P06213-2) followed at its C terminus by the 25-residue sequence SSGPSGSHHHHHHHHGSLEVLFQGP (i.e. a protease-resistant linker, the 8xHis tag, and the human rhinovirus 3C protease cleavage site) and a tandem-affinity purification tag (Rigaut et al., 1999) was cloned into the pTT6 vector, called pTT6-IRA.ECD-8xHis-TAP, for transient expression in mammalian cells. The pTT6 vector, which was derived from pTT3 (Durocher et al., 2002), featuring a Kozak sequence and a modified multiple cloning site, was kindly provided by the Protein Expression Purification and Characterization facility at the Max Planck Institute of Molecular Cell Biology and Genetics, Dresden, Germany (MPI-CBG). FreeStyle HEK293F cells (Invitrogen/Thermo Fisher Scientific, Cat# R79007, RRID:CVCL_D603) were maintained in suspension in protein-free chemically-defined FreeStyle 293 Expression Medium (Invitrogen / Thermo Fisher Scientific, Cat# R79007) supplemented with 1x Penicillin/ Streptavidin (Thermo Fisher Scientific, Cat# 15140122) at 90 rpm, 8% CO_2_, 37 °C. Prior to transfection, the medium was replaced with fresh antibiotic-free medium. 2 liters FreeStyle HEK293F cells were transiently transfected with pTT6-IRA.ECD-8xHis-TAP at a density of 2 x 10^6^ cells/ml by transfection with 2 mg endotoxin-free DNA pre-complexed with polyethylenimine (at a ratio of 5:1 w/w to DNA,) (Longo et al., 2013). Upon transfection, cells were maintained for 64 h at 31 °C, 8% CO_2_, 90 rpm. The conditioned medium was harvested by pelleting the cells at 300 x g, 10 min, 25 °C. Cells could be maintained for three more days in fresh medium for a second round of purification.

### Affinity purification of IR-ECD

IR-ECD was purified from a 2-liter batch of conditioned medium. The medium was cleared by centrifugation at 2,500 x g, 10 min, 4 °C and the supernatant was then allowed to bind to 4 ml IgG Sepharose beads for 3 h at 4 °C under constant agitation and subsequently loaded onto a 2.5 x 20 cm Econo-Column glass chromatography column (#7372522, Bio-Rad Laboratories). The flow-through was collected and then re-loaded onto the column. Running buffers were all based on HEPES-buffered saline (HBS) (50 mM HEPES, pH 7.5, 150 mM NaCl) and all purification steps were carried out at 4 °C. The IgG Sepharose beads were washed with 10 column volumes (CV) running buffer (RB) (150 mM NaCl, 50 mM HEPES, pH 7.5, 5% (v/v) glycerol), 2 CV RB-ATP (RB + 5 mM ATP, 10 mM MgCl_2_), 2 CV RB-EDTA (RB + 20 mM EDTA) and then 20 CV elution buffer (RB + 15% (v/v) glycerol). For elution IgG beads were incubated with glutathione S-transferase (GST)-tagged human rhinovirus 3C protease (50 μg protease per ml beads; provided by the MPI-CBG) for TAP tag cleavage overnight at 4 °C. After protease cleavage, IR-ECD was eluted in one step with 2.5 CV elution buffer. To remove co-eluting protease and other impurities, the eluate was incubated with 1 ml Ni-NTA Superflow beads (Quiagen, cat# 30430) for 3 h at 4 °C on a rotating wheel for immobilized metal ion affinity chromatography (IMAC). The slurry was then loaded onto a disposable conical 0.8 x 4 cm polypropylene column (Bio-Rad Laboratories). The flow-through was collected and re-loaded onto the column. The resin was washed with 10 CV HBS-20 (HBS with 10 mM imidazole (Carl Roth, Karlsruhe, Germany, Cat# 3899.3) and eluted with HBS + 280 mM imidazole in 1 CV fractions. The pH of the wash and elution buffers was adjusted to 7.5. Elution fractions were analyzed by reducing SDS-PAGE using precast NuPAGE 4-12% Bis-Tris gels (Thermo Fisher Scientific) with 1x MOPS buffer (Thermo Fisher Scientific) and subsequent Coomassie Brilliant Blue G-250 staining (Fig. S1). The most concentrated IR-ECD-containing fraction was either desalted using disposable 8.3 ml Sephadex G-25 PD-10 desalting columns (GE Healthcare) equilibrated with HBS by centrifugation, according to the supplier’s specifications, or by FPLC. The IR-ECD concentration was estimated using a molar extinction coefficient for the dimeric IR-ECD of 280,260 M^-1^ cm^-1^ at 280 nm absorbance (as calculated by ExPASy / ProtParam (Gasteiger et al., 2003) assuming one free thiol group per monomer (Chiacchia, 1991; Sparrow et al., 1997). This concentration estimate was confirmed once with a BCA protein assay (Pierce / Thermo Fisher Scientific). The protein was kept on ice until analysis. IR-ECD was characterized by analytical gel filtration using a Superose 6 Increase 5/150 GL column equilibrated with HBS with a flow rate of 0.15 ml/min at room temperature. The apparent molecular weight was estimated in SDS-PAGE with 3-8% Tris-Acetate gels (Life technologies) using HiMark unstained protein standards (Thermo Fisher Scientific).

### Microscale thermophoresis to determine insulin binding to IR-ECD

Recombinant human insulin was purchased from Sigma-Aldrich (Cat# I2643, LOT SLBR9404V, expressed in yeast, 99% purity (HPLC), 0.4% Zinc) and resuspended in 5 mM HCl at 3 mg/ml (Cat# 20252.244, VWR Chemicals, Dresden, Germany). The purity of insulin was confirmed by mass spectrometry under denaturing conditions where only monomeric and dimeric insulin with a mass of 5803.651 +/-0.003 Da was detectable, corresponding to the expected mass of insulin with all 3 disulfide bridges formed. Under native conditions, as expected, additional peaks corresponding to higher insulin oligomers appeared.

Insulin binding to the IR-ECD was analyzed by microscale thermophoresis (MST). Immediately after IMAC elution, IR-ECD was directly applied to a Superdex 200 Increase 300/10 GL column (GE Healthcare) at 0.5 ml/min in HBS at room temperature and the two peak fractions were pooled for ligand binding assays. IR-ECD was diluted to a final concentration of 100 nM in HBS-T (HBS, pH 7.5, 0.05% Tween-20) and labelled with 25 nM tris-nitriloacetic acid conjugated to NT647 (RED-tris-NTA) (Lata et al., 2005; Bartoschik et al., 2018), which was a kind gift of Jacob Piehler (University Osnabrück, Germany). A dye-to-protein molar ratio of 1:4 was chosen to circumvent interference of free dye. The reaction was incubated for 30 min in the dark at room temperature and was subsequently centrifuged at 14,000 x g for 10 min at 4 °C.

For ligand binding assays, a 100 μM stock solution of recombinant human insulin in 5 mM HCl was diluted to a concentration of 5 μM in HBS-T. A serial dilution was prepared with ligand binding buffer (i.e. HBS with 25 μM HCl). 10 μl of the diluted ligand was incubated with 10 μl of 20 nM IR-ECD overnight at 4 °C. Thus, the final assay concentrations were 10 nM IR-ECD and 2.5 nM RED-tris-NTA.

Microscale thermophoresis was carried out in standard capillaries (Cat# MO-K022, Nanotemper Technologies, Munich, Germany) on a Monolith NT.115^Pico^ instrument (Nanotemper Technologies) using the Pico-RED detector with 30% LED power at 25 °C. Data were analyzed with MO.Affinity Analysis 2.2.7 software (NanoTemper Technologies). The integral of thermophoresis traces between 1 s to 20 s on-time was used for EC50 determination and the normalized flourescence difference ΔF_norm_ was plotted against ligand concentration for dose-response plots. A concentration range was determined within which insulin did not appear to interact nonspecifically with the labelling reaction or with the dye itself, by titrating insulin against a RED-tris-NTA-labelled control peptide. A peptide comprising an 8xHis tag and the HRV3C cleavage site (i.e. HHHHHHHHLEVQL) served as control peptide.

### Thermal stability

To further characterize IR-ECD and to monitor its stability, a thermal unfolding assay was carried out applying label-free low-volume differential scanning fluorimetry (nanoDSF). IR-ECD was diluted to 500 nM in HBS and incubated with or without 50 μM insulin for 1 h on ice in a volume of 22 μl. Samples were loaded into nanoDSF Grade Standard capillaries (Cat# PR-C002, NanoTemper Technologies) in duplicates and transferred to a Prometheus NT.48 instrument (NanoTemper Technologies). Thermal unfolding was detected by recording the intrinsic tryptophan fluorescence (emission ratio at 350 nm and 330 nm) during heating in a linear thermal ramp (1 °C/min; 20 °C to 95 °C) with an excitation power of 100%.

### Cryo-EM grid preparation and imaging

The human IR-ECD was purified as described above, desalted after immobilized metal affinity chromatography elution using disposable 8.3 ml Sephadex G-25 PD-10 desalting columns, concentrated to 3 μM in Amicon Ultra-0.5 ml ultrafiltration units with ultracel-100 membranes (Merck Chemicals), and stored at 4 °C until further use. Prior to cryo-EM sample preparation, the concentrated protein was gel-filtrated using a Superdex 200 10/300 GL column equilibrated in HBS at a flow rate of 0.5 ml/min and 4 °C (GE Healthcare). Peak fractions at a final OD_280nm_ of ∼ 0.4 (∼ 1.4 μM) were incubated for ∼ 30 min at 4 °C with or without recombinant human insulin supplementation at 35x molar excess (∼ 50 μM final concentration). 4 μl of these samples were applied to glow discharged (2.2×10^-1^ mbar for 2×20sec) Quantifoil holey carbon grids (R2/1, 200 mesh, Quantifoil). The grids were plunge vitrified into a liquid ethane/propane mix using a Vitrobot Mark IV at 4 °C and 95% humidity. Cryo-EM data were collected on a FEI Titan Krios microscope operated at 300 kV, equipped with a post-column GIF and a K2 Summit direct detector operating in counting mode. A total of 8882 movies were recorded at a nominal magnification of 130,000x that corresponds to 1.059 Å/pixel at the specimen level using SerialEM (Mastronarde, 2005). The total exposure of 55 e^-^/Å^2^ at the specimen level was evenly distributed over 51 frames during 10.2 s. The preset target defocus range was 0.5 to 3.5 μm. The sample preparation and data collection strategies for the *apo*-IR-ECD samples were very similar except that no insulin was used for grid preparation. These data were collected with a total exposure of 59 e^-^/Å^2^ spread over 51 frames and 10.2 sec. The target defocus ranged between 0.5 and 3.5 μm. No stage pretilt was employed for either of the two data sets.

### Cryo-EM data acquisition and processing

The RELION-3.0 implementation of MotionCor2 (Zheng et al., 2017) was used to correct for beam-induced sample motions and radiation damage. The summed and dose-weighted micrographs were used for further processing. Particles were selected using Gautomatch (http://www.mrc-lmb.cam.ac.uk/kzhang/Gautomatch/). CTF parameters were determined using Gctf (Zhang, 2016). If not stated otherwise all further processing was carried out in RELION-2.1 or RELION-3.0 (Kimanius et al., 2016; Zivanov et al., 2018). In the case of the insulin-bound structure initial analysis of particles picked without templates yielded a 3D reconstruction using as template a 60 Å low-pass filtered initial model generated by the stochastic gradient descent implementation of RELION-2.1 (Kimanius et al., 2016) (cp. Fig. S4 for a graphical overview of the processing routine). Low pass filtered projections of this reconstruction were used as templates for template-based particle picking on all micrographs. This resulted in 2 997 079 particle candidates. The particle stack was cleaned up by unsupervised 2D classification in subsets of ∼ 120 000 particles. Subsequently, the data were further processed in ∼ 120 000 particle chunks in 3D classification with the first reconstruction as a 60 Å low-pass filtered starting model. The resulting cleaned data set of 326 257 particles reached a nominal global resolution of 4.9 Å after 3D-refinement and postprocessing. Bayesian polishing in RELION-3.0 (Zivanov et al., 2018) was employed to correct further for beam induced motion and radiation damage, improving the quality of the map to a final apparent resolution of 4.3 Å. The global resolution estimates of the obtained reconstructions are quoted as good proxies for the overall relative quality of the individual reconstructions, fully acknowledging the differences in local resolution estimates as well as the anisotropy of the data. The angular distribution of particles contributing to this map is shown in Fig. S5C and S5D, the Fourier shell correlation (FSC) curve of the masked independent half-maps in Fig. S5E. The rotation vs. tilt angle plot in Fig. S5C was created by binning the angular assignments of all particles contributing to this reconstruction in 3° by 1.5° bins, respectively, followed by plotting the resulting distribution using the tidyverse collection of R packages. The local resolution estimate in Fig. S5F was calculated with the local resolution routine implemented in RELION-3.0 (Zivanov et al., 2018).

The data of the ligand-free IR-ECD sample were processed using a similar approach as outlined above. Since the attempts at 3D reconstruction never yielded resolution in the sub-nanometer range, only 2D class averages are shown.

### Model Building and Refinement

A non-glycosylated IR-ECD model complexed by 4 insulins was constructed initially for fitting into the cryo-EM density map. This model was based on the previously published, partial, insulin-bound IR-ECD (PDB:6CEB, including the two head-bound insulins 1 and 1’). The L1 and L1’ domain residues H144 were modified to Y144 to match the IR construct used here (uniprot entry P06213-2). Regions that were not resolved in PDB:6CEB were added as described in the following. As reliable starting models for the FnIII domains, we included the respective coordinates from PDB:4ZXB (Croll et al., 2016) in the structure. Additionally, we constructed tentative models of the IDα loops (chain α and α’ residues 651-687) using MODELLER (Eswar et al., 2006) and included them in the structure since we observed some incohesive density features for these regions. The stalk-bound insulins 2 and 2’ were modeled based on the structure of porcine insulin (PDB:4INS) (Baker et al., 1988). To match the human insulin sequence, the insulin B-chain C terminus residue was mutated from A30 to T30. The model is thus consistent with the complete human IR-ECD (UniProt ID: P06213-2) and human insulin sequences (UniProt ID: P01308), and matches the experimental constructs used in this study.

As a first step in the fitting procedure, global, rigid body docking of the resulting non-glycosylated IR-ECD in complex with 4 insulins into the density map was performed in UCSF Chimera (Pettersen et al., 2004). To locally improve the model we employed a combination of flexible fitting methods including the real-space structure refinement program DireX (Wang and Schroder, 2012), followed by the simple relax protocol in torsional space in Rosetta (Fleishman et al., 2011; Conway et al., 2014). As a last step, the cysteines involved in intra- and inter-chain dimer bonds as well as specific β-strands of the FnIII-3 domains were directed into selected regions of the density map by interactive molecular dynamics flexible fitting (iMDFF) (Trabuco et al., 2009; McGreevy et al., 2016).

After completion of the initial fitting routine outlined above, the structure was subjected to several rounds of iterative real-space refinement in phenix.refine (Afonine et al., 2018) and manual adjustment in Coot (Emsley et al., 2010). Progress in modelling was monitored via the map-to-model correlation coefficients, geometry indicators, and the map-versus-model FSC (see Tab. S1). Structure images were created in PyMOL2 (PyMOL Molecular Graphics System, Schrödinger, LLC.) and ChimeraX (Goddard et al., 2018). The refined model will be deposited in the PDB and is referred to in the main text as “cryo-EM structure”. It is displayed graphically in all structure figures except Fig. S7.

Since our reconstruction did not produce clear density features for the Arg-Lys-Arg-Arg residues (furin cleavage site), the disordered IDβ region and the C-terminal purification tag sequence, these parts were not included in the refined structure (see also Tab. S2). However, for completeness, they were included in the model used in molecular dynamics simulations described below.

### Atomistic molecular dynamics (MD) simulations

For atomistic MD simulations, we completed the structure refined against the EM map described above by adding all the loops invisible in our density map, i.e. the furin cleavage site (residues 720-723), the disordered and highly glycosylated N-terminal region of the IRβ subunit (residues 724-756), and the C-terminal His-tag sequence). All of these additional loops and regions were built using MODELLER (Eswar et al., 2006). Additionally, we added 17 N-linked and 6 O-linked glycans on each monomer using the doGlycans tool (Danne et al., 2017) based on the glycan composition defined by Sparrow et al. (2007) and Sparrow et al. (2008) (Tab. S3). The OPLS-AA force field (Kaminski et al., 2001; Danne et al., 2017) was used for the protein, glycans, and ions. The glycosylated IR-ECD was energy-minimized in vacuum using the steepest descent algorithm to remove any steric clashes due to overlapping atoms. The minimized structure was then solvated using the TIP3P water model (Jorgensen et al., 1983) in a box of [21 nm]^3^. The solvated structure was neutralized with an appropriate number of Na^+^ counterions complemented by 150 mM of NaCl to match experimental buffer and salt concentration. The system consisted of 924,775 atoms in total. The resulting structural model of the IR-ECD is referred to as “MD model” in the text.

Prior to MD simulations, the system was again subjected to energy minimization followed by 50-ns equilibration under NVT (constant particle number, volume, and temperature) conditions at 298 K using the v-rescale thermostat (Bussi et al., 2007) with a time constant of 0.1 ps. At this stage the IR-ECD and the insulin backbone atoms were position-restrained with a force constant of 1,000 kJ mol^-1^nm^-2^. Next, equilibration of the system was continued under NpT (constant particle number, pressure, and temperature) conditions using isotropic Parrinello-Rahman pressure coupling (Parrinello, 1981; Parrinello and Rahman, 1980, 1982) with a time constant of 2 ps over a period of 50 ns, with reference pressure set to 1 bar and isothermal compressibility to 4.5 x 10^5^ bar^-1^. The IR-ECD and the insulin backbone atoms were again position-restrained with a force constant of 500 kJ mol^-1^nm^-2^. Electrostatic interactions were calculated by the Particle Mesh Ewald method (Darden et al., 1993; Essmann et al., 1995) using 1.0 nm for the cut-off of the real space component. The same cut-off distance was set for van der Waals interactions together with the LINCS algorithm (Hess et al., 1997) for all bonds. Periodic boundary conditions were applied in all three dimensions. The final production run for 500 ns was carried out after removal of all position restraints, and the rest of the input parameters were the same as those used under NpT equilibration simulations. All MD simulations were carried out with an integration time step of 2 fs using the GROMACS 4.6 simulation package (Hess et al., 2008) and the output trajectory and energies were saved every 100 ps. For reproducibility of the results, 10 repeats (500 ns each) were performed.

Analyses made use of GROMACS tools, in-house built scripts, and contact maps built with the g_distMat analysis tool. A contact for a given pair of residues was considered to be established if the minimum distance between any atoms in the two residues was ≤ 3.5 Å or ≤ 6 Å. Root-mean-square deviation (RMSD) analysis was performed for backbone atoms with respect to the starting MD model. The final 100 ns from each of the 10 trajectories were used to generate residue contact occupancy maps. MD movies and figures were prepared using VMD (Humphrey et al., 1996) and PyMOL2.

## Data availability

The cryo-EM density maps and the atomic model developed in this study will be available from the Electron Microscopy Data Bank and Protein Data Bank, respectively, after peer-reviewed publication.

## Conflict of interest

The authors declare no conflict of interest.

## Acknowledgments

We would like to acknowledge Elena Conti (MPIB) for generous and unconditional support. Generous computational resources were made available by the High-Performance Computing Center of the TU Dresden (UC), the Leibniz Supercomputing Centre of the Bavarian Academy of Sciences and Humanities (UC; project pr48ci) and the CSC-IT Centre for Science Espoo, Finland (IV). Financial support was provided by: the Academy of Finland (Center of Excellence program) (IV); the European Research Council (Advanced Grant CROWDED-PRO-LIPIDS) (IV), the Deutsche Forschungsgemeinschaft (DFG, German Research Foundation) – Project Number 251981924 – TRR 83 (ÜC, IV), Project Number 347368302 – FOR2682 (UC, TG) and the Dresden International Graduate School for Biomedicine and Bioengineering – Project Number 24185540 (TG). Cryo-electron microscopy experiments were carried out at the cryoEM Facility at the MPIB (Martinsried).

