## Supplementary Information for "Cryo-EM structure of the complete and ligand-saturated insulin receptor ectodomain"

**Running title: Molecular basis for site 2 insulin-insulin receptor interaction**

Gutmann, Theresia[1,2#]; Schäfer, Ingmar[3#]; Poojari, Chetan[4#]; Brankatschk, Beate[1,2]; Vattulainen, Ilpo[4,5*]; Strauss, Mike[6*]; Coskun, Ünal[1,2*]

[1]Paul Langerhans Institute Dresden of the Helmholtz Zentrum Munich at the University Hospital and Faculty of Medicine Carl Gustav Carus of TU Dresden, Dresden, Germany

[2]German Center for Diabetes Research (DZD e.V.), Neuherberg, Germany

[3]Department of Structural Cell Biology, Max Planck Institute of Biochemistry, Munich, Germany

[4]Department of Physics, University of Helsinki, P.O. Box 64, FI-00014, Helsinki, Finland

[5]Computational Physics Laboratory, Tampere University, Tampere, Finland

[6]Department of Anatomy & Cell Biology, McGill University, Montreal, Quebec, Canada

### these authors contributed equally

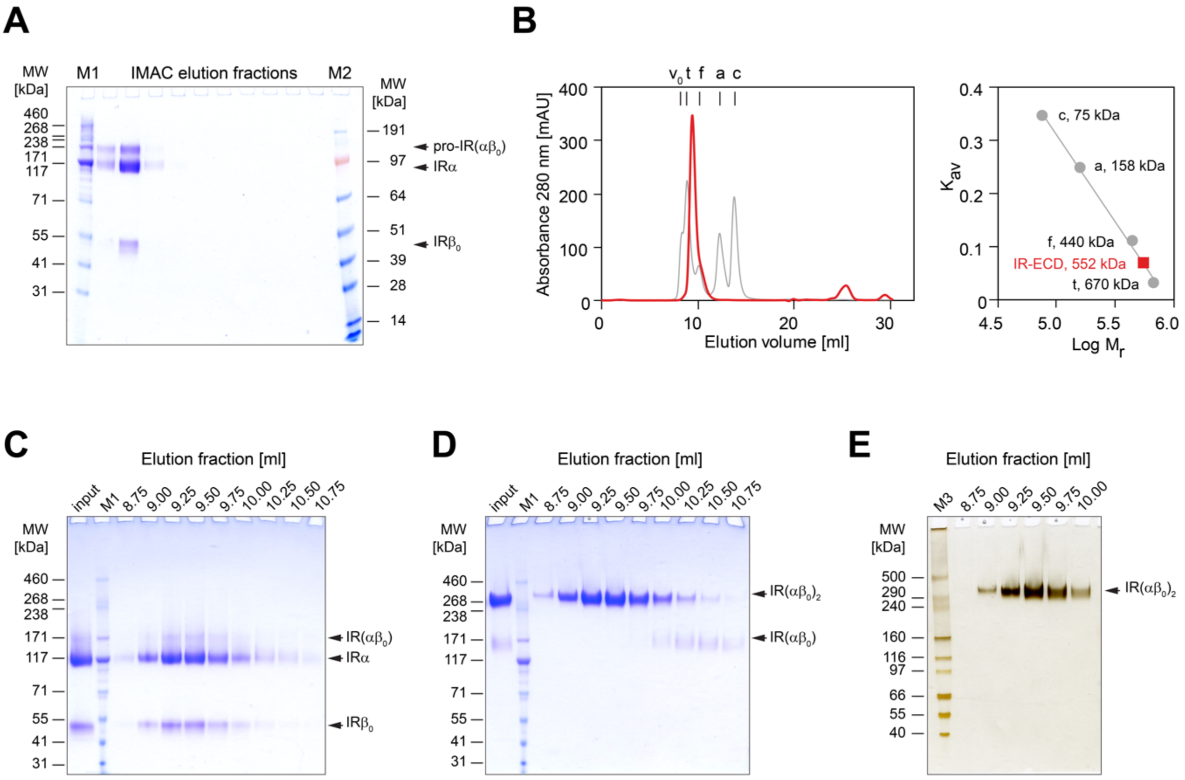

**Figure S1: Purification of IR-ECD.** (**A**) Coomassie G-250 BrilliantBlue-stained 4-12% Tris-Bis gel run in MOPS buffer of the IMAC elution fractions under reducing conditions. (**B**) The peak fraction containing IR-ECD was gel-filtrated through a Superdex 200 Increase 10/300 GL column. The void volume ($v_0$) and elution volumes of the standards bovine thyroid thyroglobulin (t), horse spleen ferritin (f), rabbit muscle aldolase (a), and egg white conalbumin are indicated. The partition coefficient ($K_{av}$) is plotted against the logarithm of molecular weight for standards (right) to determine the IR-ECD apparent molecular weight, which is with 552 kDa is considerably larger than in denaturing SDS-PAGE, due to its elongated shape. Samples of eluted fractions were analysed by SDS-PAGE on 3–8% Tris-Acetate gels under (**C**) reducing and (**D**) non-reducing conditions and gels were subsequently stained with Coomassie G-250 BrilliantBlue. (**E**) Complete silver-stained SDS-PAGE gel corresponding to the single lane shown in Fig. 1B. The apparent molecular weight was estimated to 351 kDa for the dimeric IR-ECD ($IR(\alpha\beta)_0$), 120-130 kDa for the alpha subunit (IRα), and 50–54 kDa for the extracellular IR beta ($IR\beta_0$) subunit, based on HiMark unstained protein standards on 3–8% Tris-Acetate gels. Markers used were HiMark pre-stained protein standard (M1), SeeBlue Plus2 pre-stained protein standard (M2), and HiMark unstained protein standard (M3).

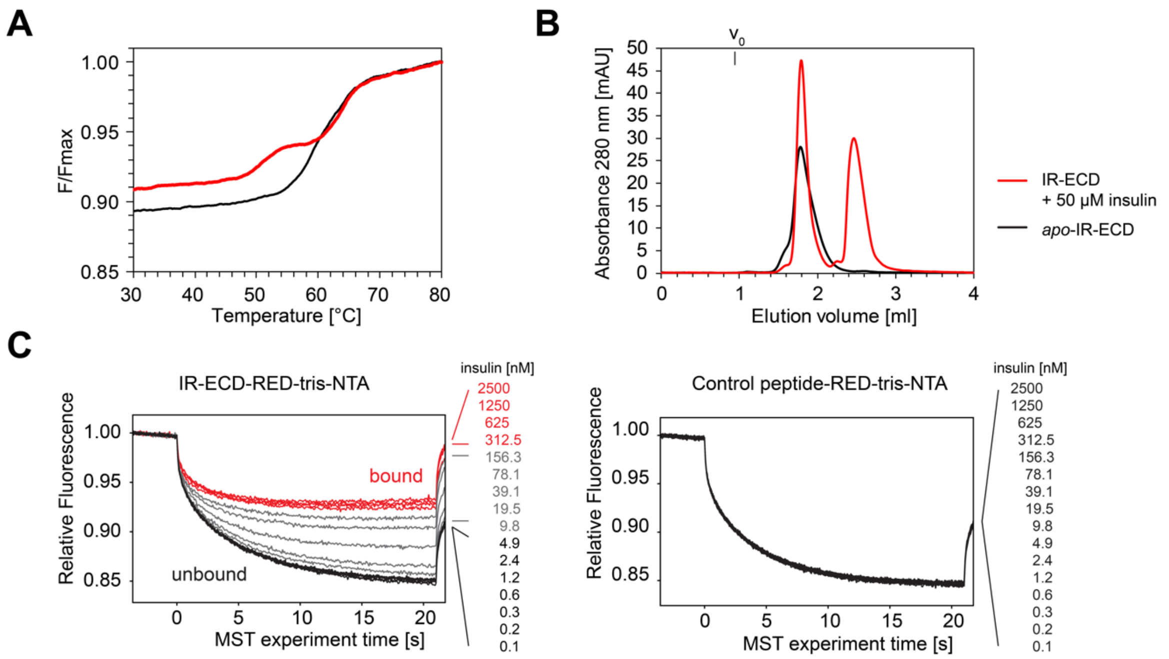

**Figure S2: Insulin binding to IR-ECD.** (**A**) The thermal unfolding of IR-ECD was assessed in the absence (black) or presence (red) of 50 µM human insulin by recording intrinsic tryptophan autofluorescence ratios at 350 nm and 330 nm. The plot shows normalized tryptophan autofluorescence over temperature. (**B**) Superose 6 size exclusion chromatograms of IR-ECD in the absence (black) or presence (red) of insulin. Compared to *apo*-ECD, insulin-bound IR-ECD elutes in a sharper peak followed by a second peak of unbound insulin. (**C**) Representative MST traces of 10 nM IR-ECD (labelled with RED-tris-NTA) after exposure to insulin. Native insulin at concentrations from 2.5 µM to 76 pM was titrated against 10 nM soluble RED-tris-NTA-labelled IR-ECD. The corresponding dose-response curve is plotted in Fig. 1C, with a half maximal effective concentration ($EC_{50}$) of 39 nM insulin. To rule out non-specific interactions or interference with the labelling strategy, MST traces of a synthetic control peptide labelled with RED-tris-NTA were recorded after exposure to insulin and confirmed not to interact with insulin at concentrations up to 2.5 µM.

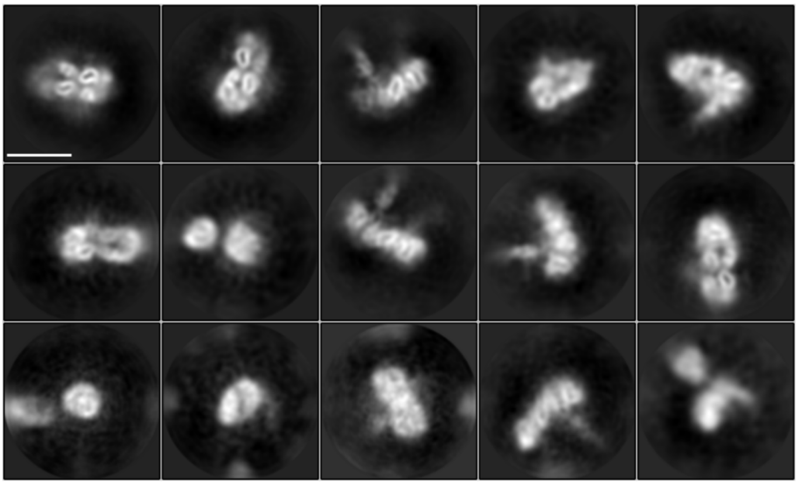

**Figure S3: 2D class averages of the *apo*-IR-ECD obtained by cryo-EM.** Scale bar 10 nm.

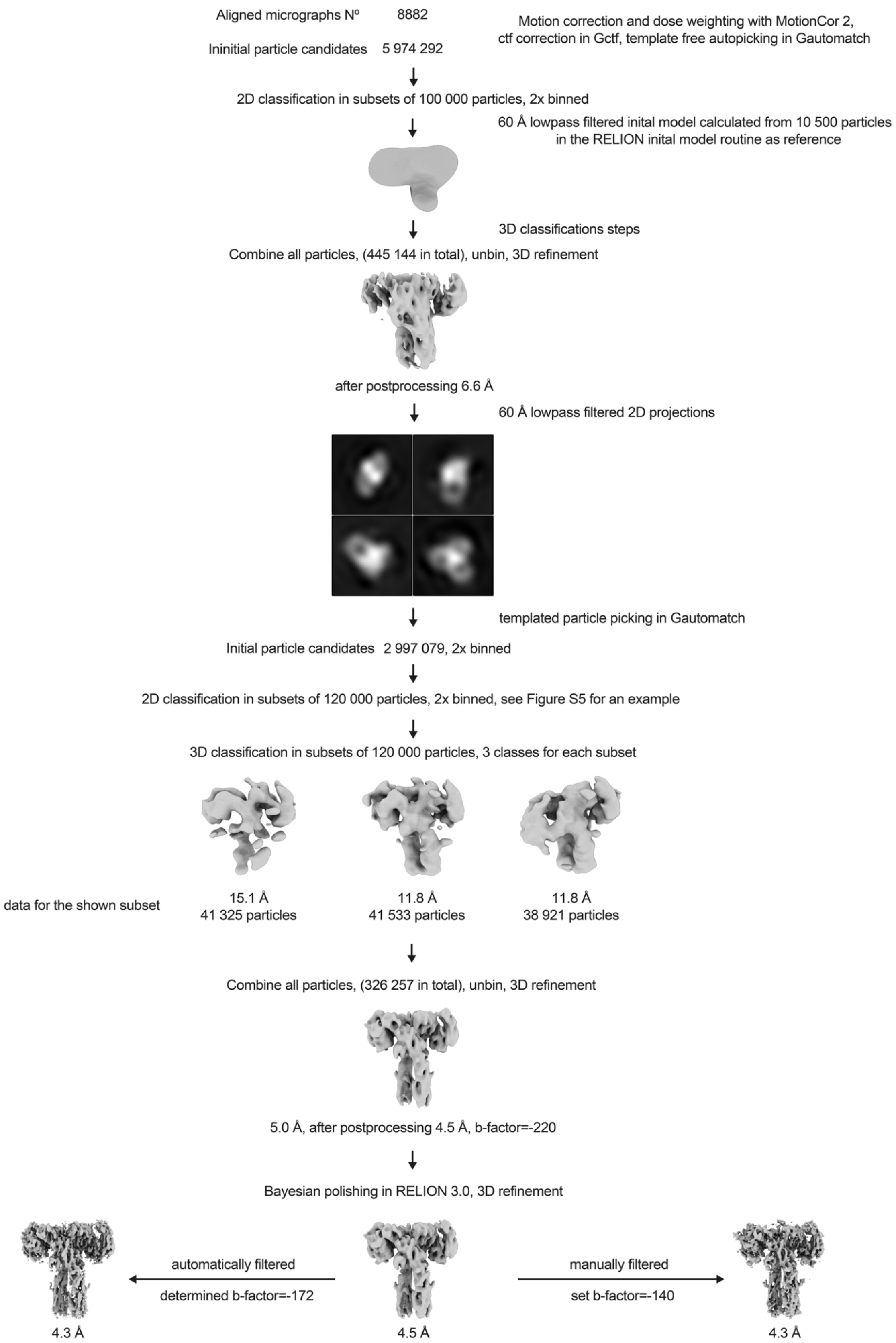

**Figure S4: Overview of the cryo-EM data processing scheme.** Particle sorting and classification scheme used for 3D reconstruction of the insulin–IR-ECD complex. The individual nominal global resolutions are quoted as good proxies for translational and rotational accuracy of reconstructions as well as for the level of detail observed in individual maps.

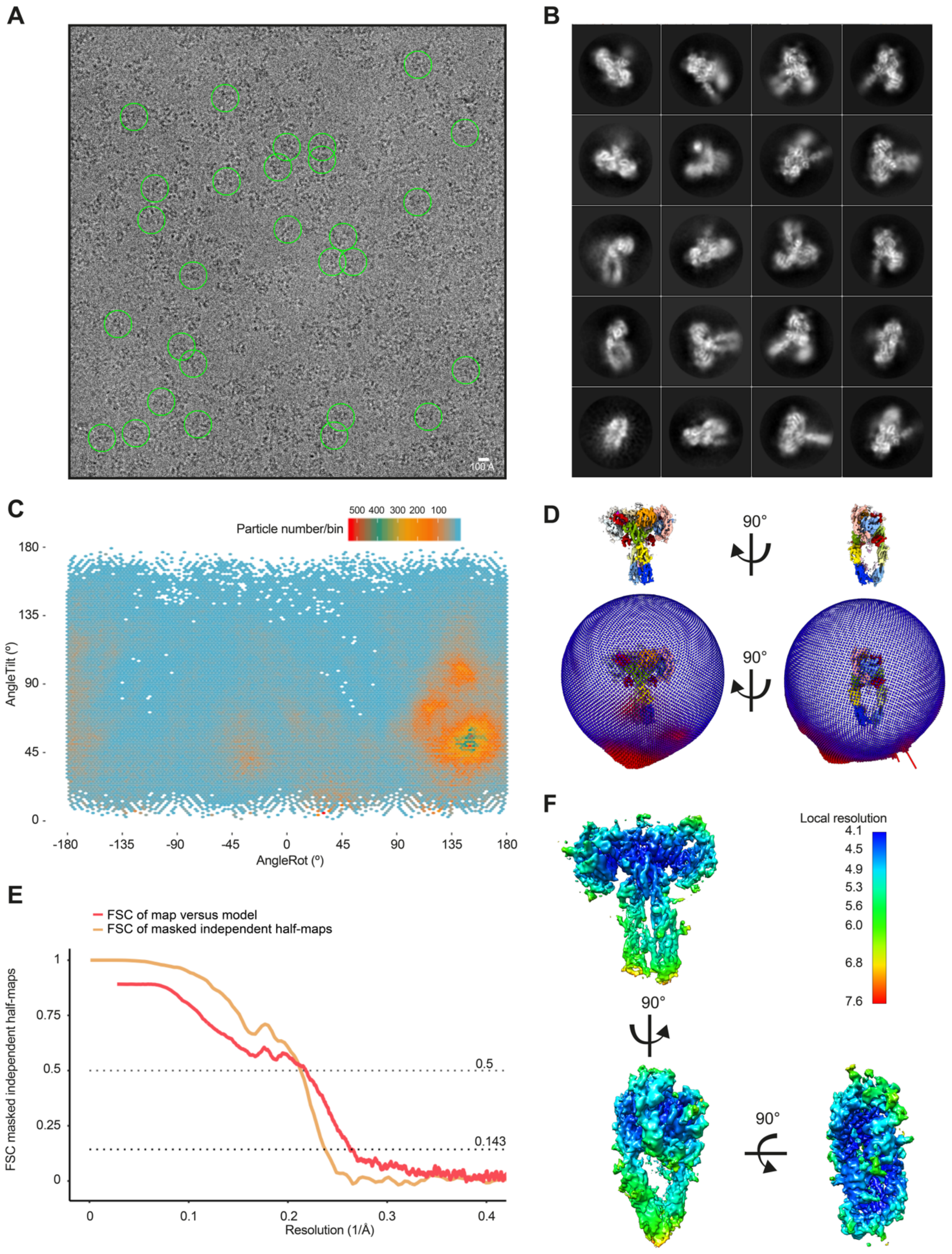

**Figure S5: Single-particle cryo-EM analysis of the insulin–IR-ECD complex.** (**A**) Representative micrographs of the insulin–IR-ECD data set. The scale bar in the cryo-EM micrograph corresponds to 100 Å and the green circles (260 Å diameter) indicate particles contributing to the final reconstruction with a nominal global resolution of 4.3 Å (see Fig. S4). (**B**) Reference-free 2D class averages of the insulin–IR-ECD complex from an initial 2D classification run (see Fig. S4 for details). Some structural heterogeneity is apparent especially in the stalk region. (**C**) Angular distribution of particles contributing to insulin–IR-ECD complex reconstruction. Tilt and rotation angles were plotted against each other for the final 4.3 Å 3D reconstruction. The color of each sampling bin indicates the number of particles in the respective bin. As in the spherical angular distribution representation in (**D**), blue denotes fewer red more particles (326 257 particles in total). (**E**) Fourier Shell Correlation (FSC) of masked independent half-maps and of map-versus-model of the final insulin–IR-ECD reconstructions used for modeling and structure interpretation (cp. Fig. S4 for details). The nominal global resolution of the full insulin–IR-ECD complex was determined to be 4.3 Å according to the 0.143 cut-off criterion (Rosenthal and Henderson, 2003). Map-to-model correlation showed agreement at the 0.5 cut-off criterion to 4.6 Å. (**F**) Map of the insulin–IR-ECD complex colored according to local resolution estimate. The central parts of the head are resolved at higher resolution whereas distal parts of the stalks are resolved at lower resolution.

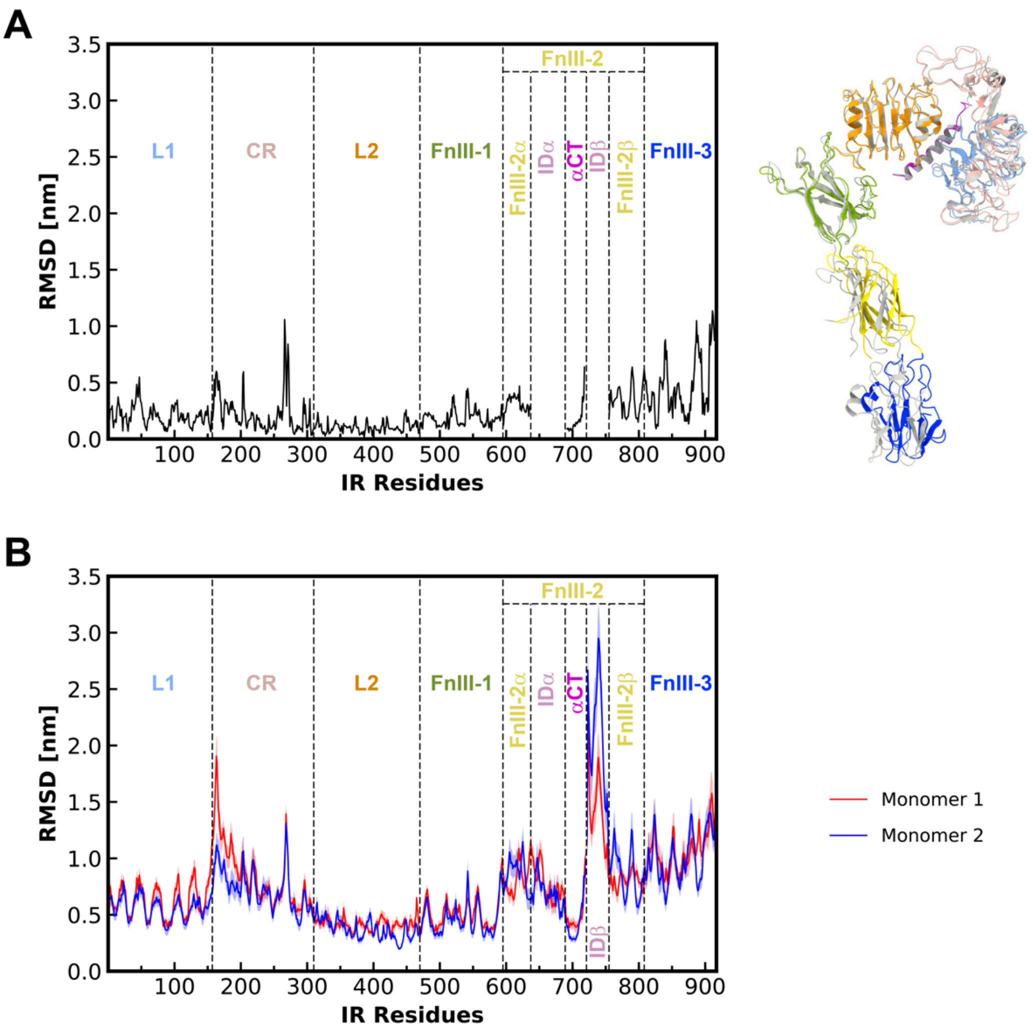

**Figure S6: Asymmetric arrangement of two IR monomers**. (**A**) The panel on the right shows the two superimposed IR monomers. Monomer 1 domains are colored as in Fig. 1 and monomer 2 is colored in grey. On the left a per-residue plot of the backbone root mean square deviation (RMSD) between the two monomers is depicted. Individual domains are separated by dashed bold lines. (**B**) Backbone RMSD measured for IR residues averaged from 10 MD simulations. Red and blue lines indicate monomer 1 and monomer 2, respectively. RMSDs were calculated over 500 ns with respect to the starting MD model.

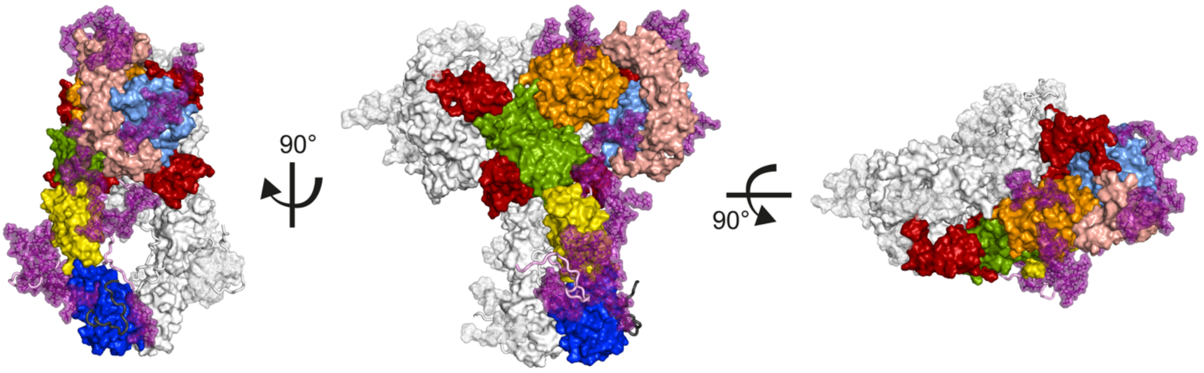

**Figure S7: Orthogonal views of the complete insulin–IR-ECD starting model used for MD simulations in surface representation.** One monomer is color-coded as in Fig. 1 with carbohydrates in purple and the insulin moieties in red. The second monomer is depicted in white. The disordered ID and C-terminal linker + tag are shown in cartoon style coloured in light violet and in dark grey, respectively.

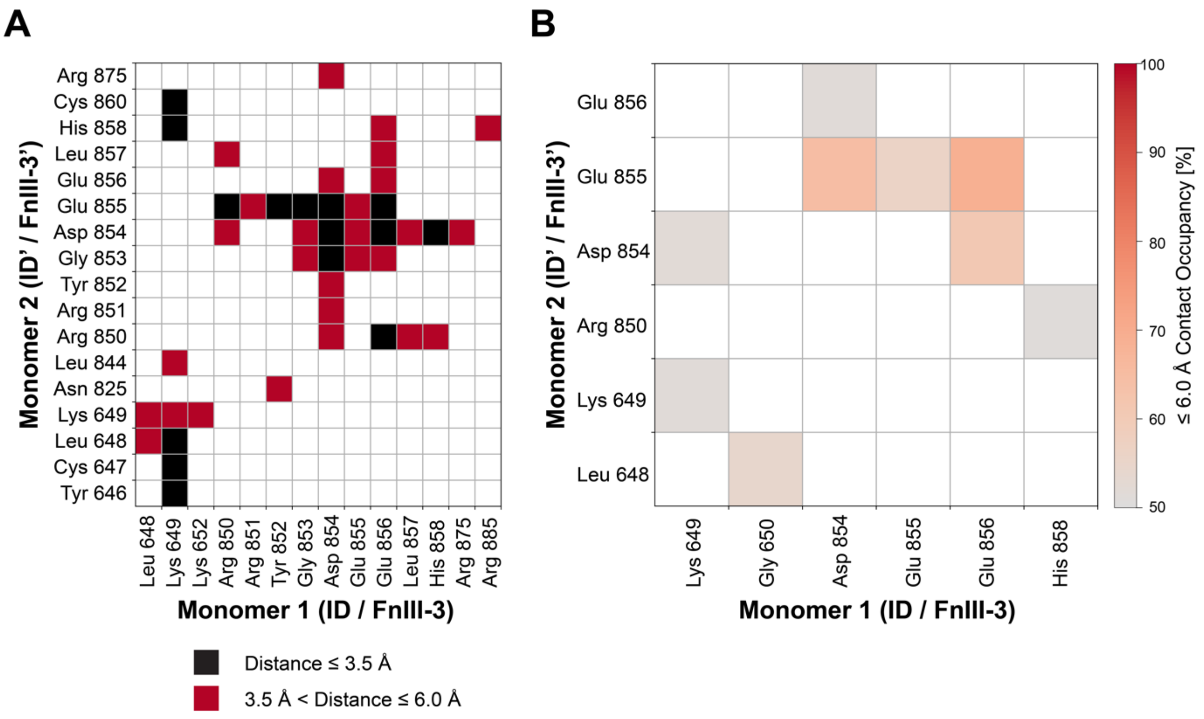

**Figure S8: Interactions of membrane-proximal FnIII-3 domains.** (**A**) Contacts calculated for the initial MD structure: Contact map showing interactions between (ID)/FnIII-3 domains from monomers 1 and 2 with contact cut-offs set to ≤ 6 Å (red) and ≤ 3.5 Å (black). (**B**) Contact occupancies calculated from MD simulations with cut-off 6 Å. Only contacts of at least 50% occupancy are displayed.

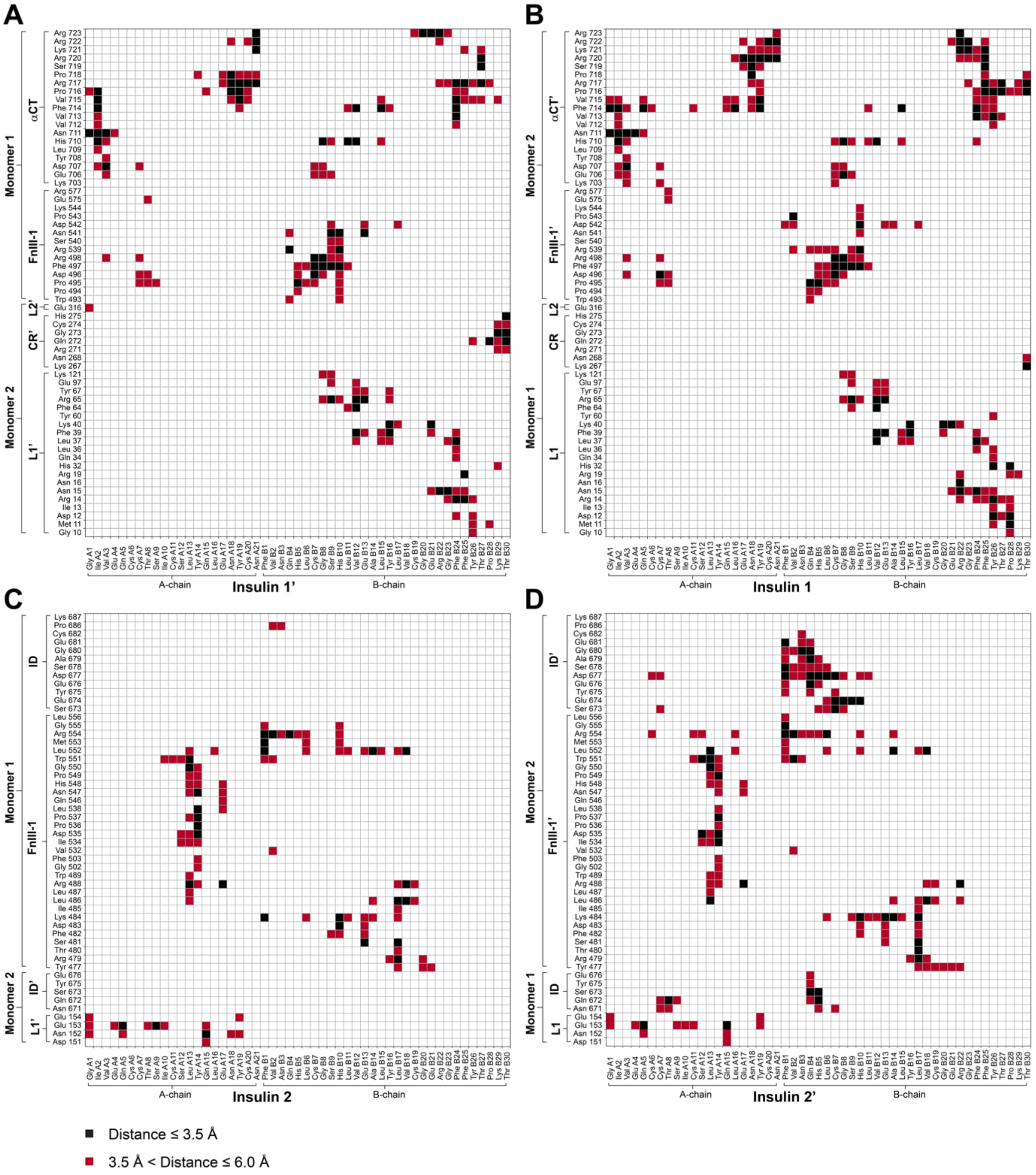

**Figure S9: Contact map for insulin–IR-ECD interactions in the cryo-EM structure with a distance cut-off of 3.5 Å and 6 Å.** Contact map showing interactions between IR-ECD and head-bound insulins 1'/1 (**A, B**) and stalk-bound insulins 2/2' (**C, D**). Contacts with a cut-off of 3.5 Å and 6 Å are shown in black and red, respectively. The arrangement of the maps corresponds to the location of the respective insulin in the ectodomain (front view).

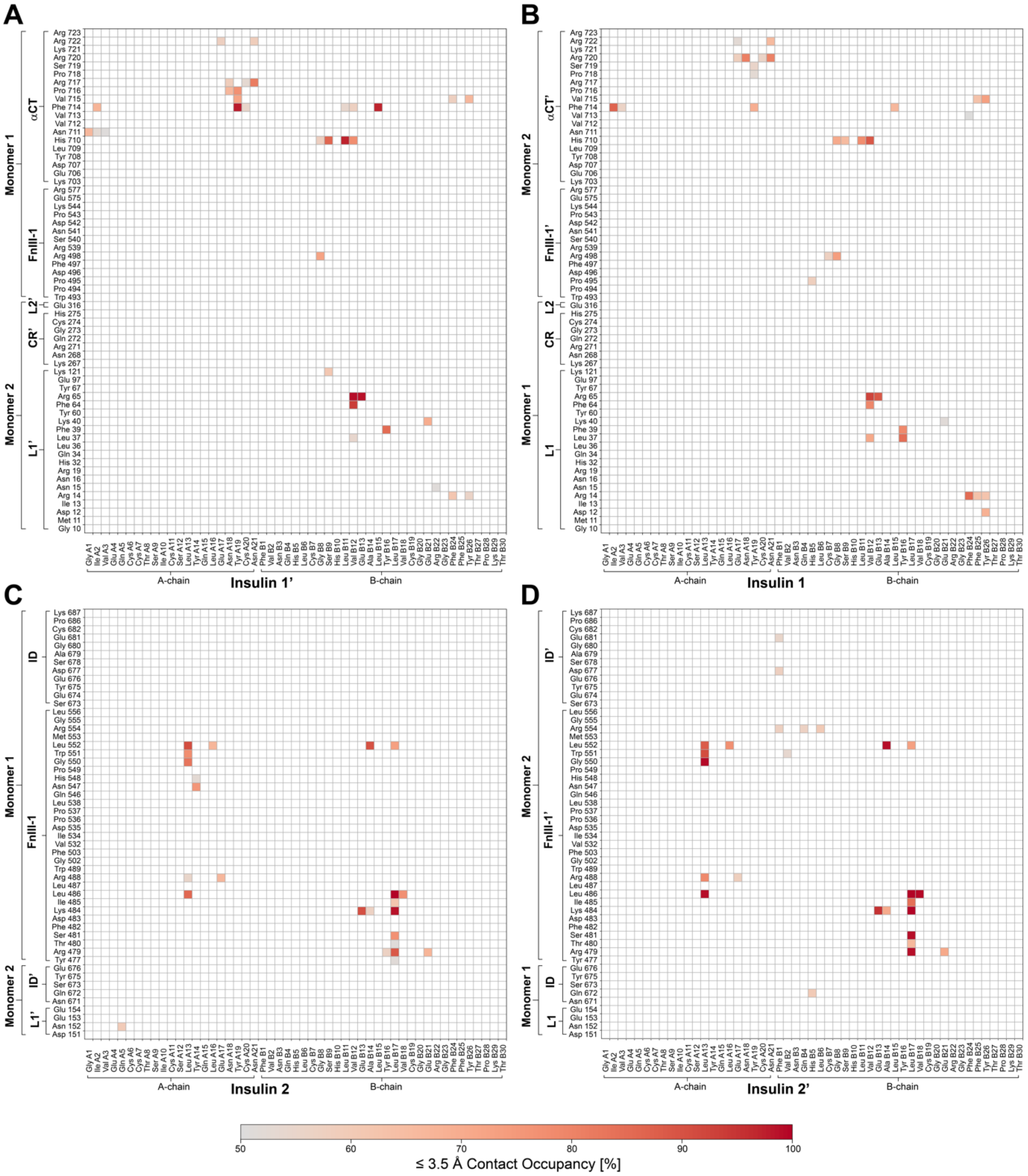

**Figure S10: Contact occupancies for insulin–IR-ECD interactions with a cut-off of 3.5 Å in the MD simulations.** Contact map showing interactions between IR-ECD and head-bound insulins 1'/1 (**A, B**) and stalk-bound insulins 2/2' (**C, D**). Only contacts with an occupancy > 50% are displayed. The arrangement of the maps corresponds to the location of the respective insulin in the ectodomain (front view).

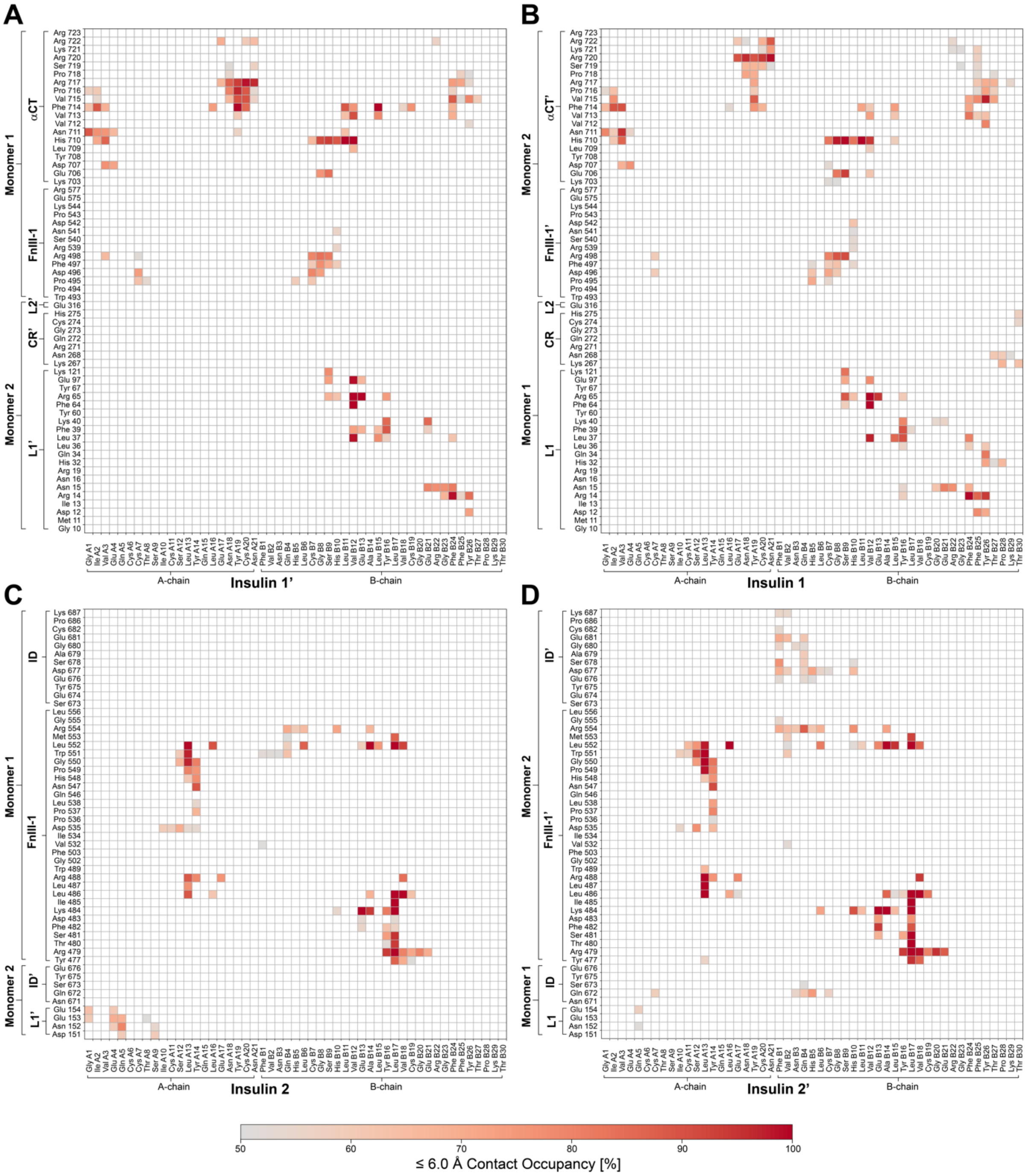

**Figure S11: Contact occupancies for insulin–IR-ECD interactions with a cut-off of 6 Å from MD simulations.** Contact map showing interactions between IR-ECD and head-bound insulins 1'/1 (**A, B**) and stalk-bound insulins 2/2' (**C, D**). Only contacts with an occupancy > 50% are displayed. The arrangement of the maps corresponds to the location of the respective insulin in the ectodomain (front view).

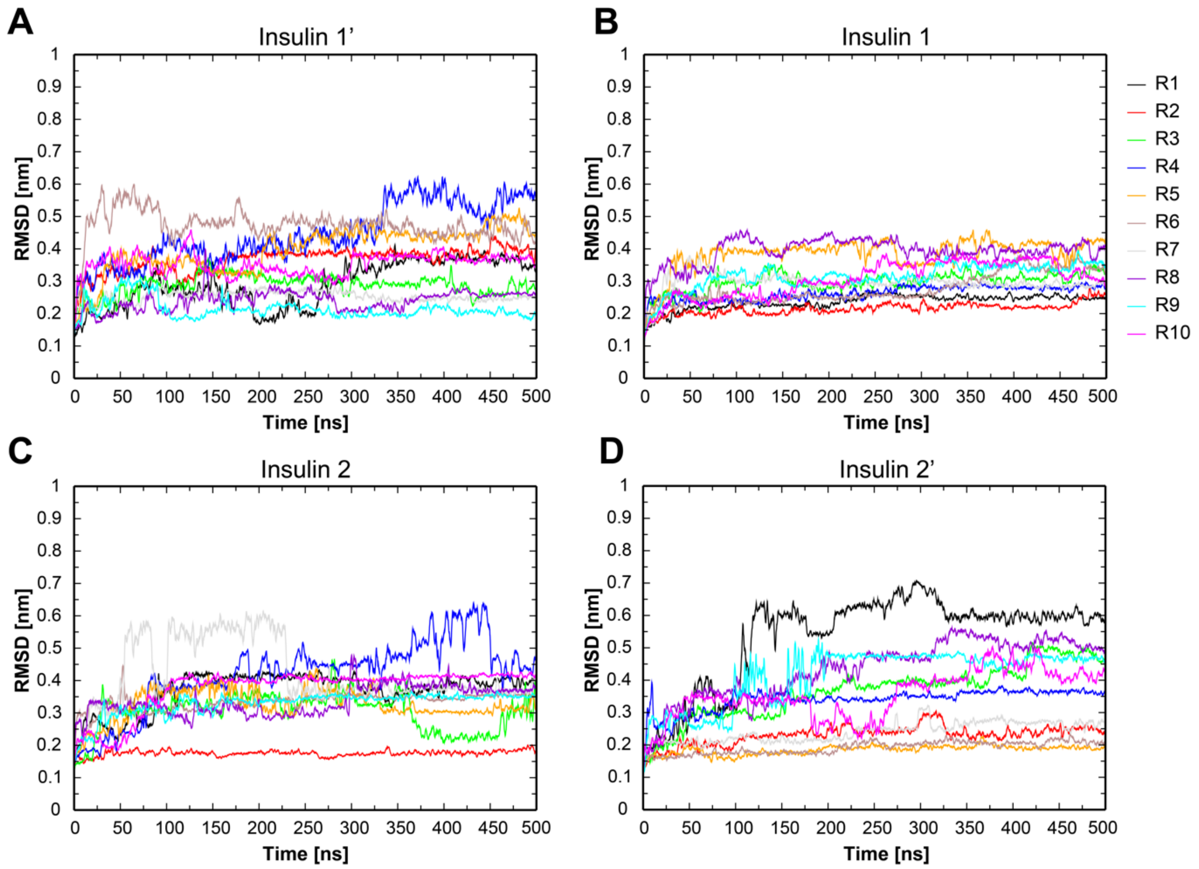

**Figure S12: Time-depended backbone RMSD for the four bound insulins determined from 10 MD simulations (R1-R10).** RMSD for head-bound insulins 1'/1 (**A, B**) and for stalk-bound insulins 2'/2 (**C, D**). RMSDs were calculated with respect to the initial MD model.

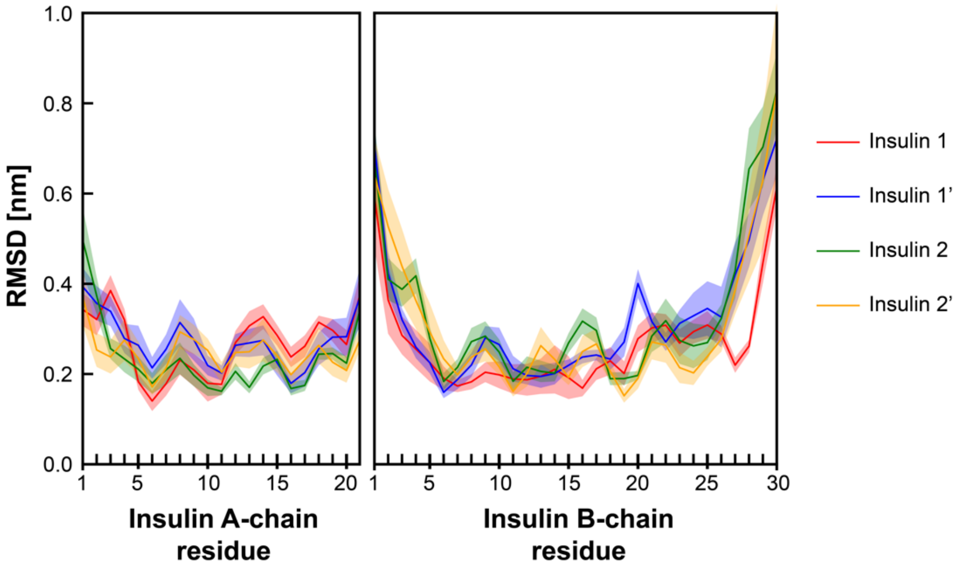

**Figure S13: RMSD determined for insulin residues in 10 MD simulations.** RMSD for insulins 1, 1', 2, and 2' averaged over 10 MD runs. The RMSDs were calculated with respect to the initial MD model.

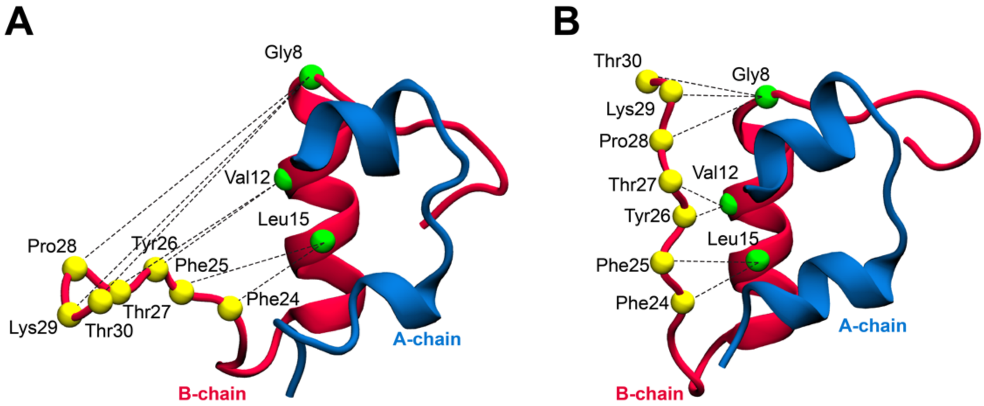

**Figure S14: Insulin B-chain C terminus dynamics.** To monitor the B-chain C terminus dynamics, we recorded the distance between residues in the B-chain alpha helix and residues within the B-chain C terminus as indicated above with dashed lines. Cryo-EM structures of insulin 1 in the open conformation (**A**) and insulin 2 in the closed conformation (**B**) are displayed in cartoon representation. See Tab. S4 for the corresponding distance measurements from the cryo-EM structure and from MD simulations.

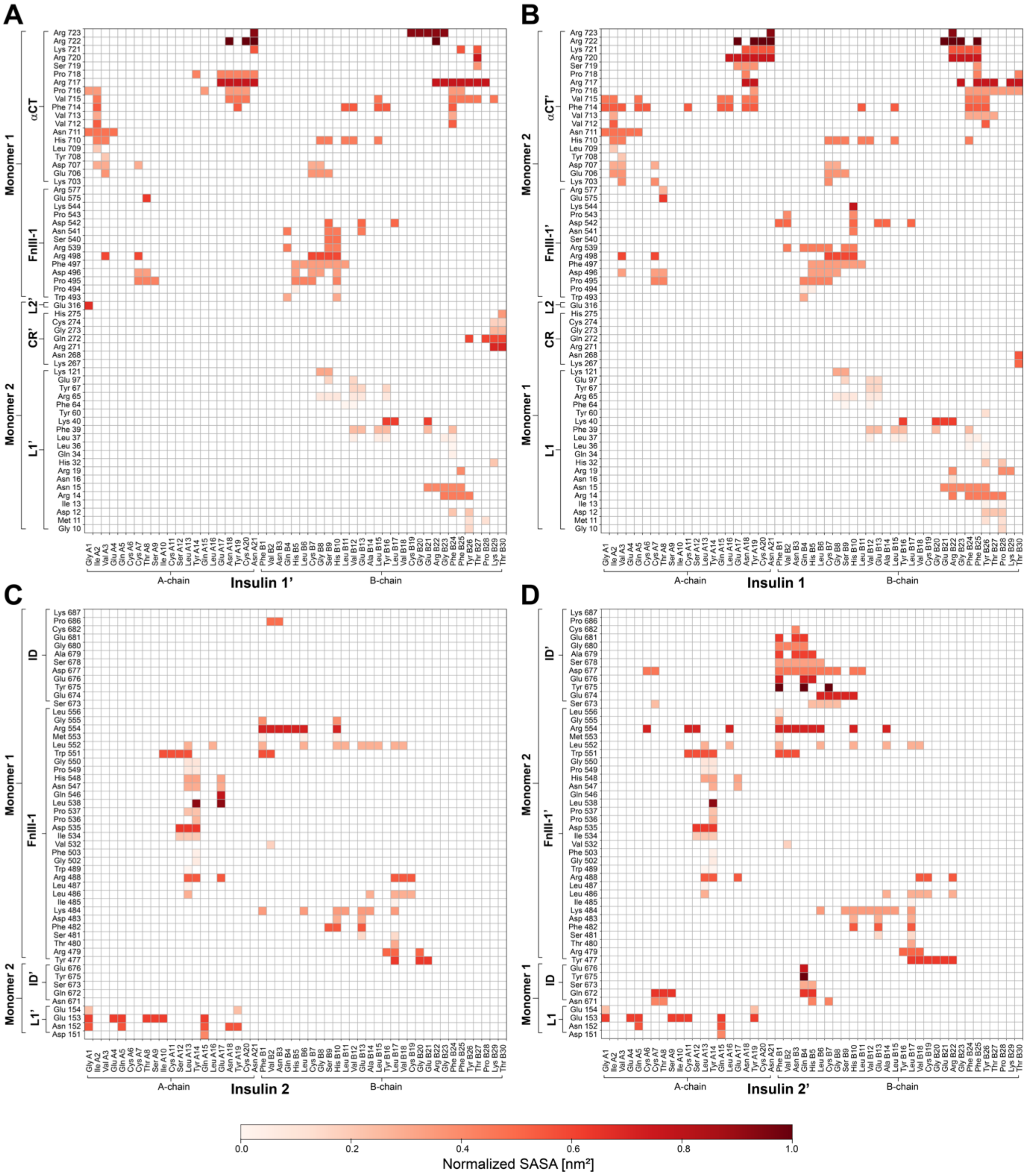

**Figure S15: Solvent-accessible surface area (SASA) for the *apo*-IR-ECD (PDB ID: 4ZXB).** SASA calculated for IR-ECD residues interacting with insulins 1'/1 (**A, B**) or insulins 2/2' (**C, D**). Interactions are based on Fig. S9 (contacts with cut-off distance < 6 Å). The arrangement of the maps corresponds to the location of the respective insulin in the ectodomain (front view).

**Table S1: Summary of cryo-EM data collection, including processing statistics and model quality indicators.**

| | |
|---|---|
| **Cryo-EM data collection** | |
| Microscope | FEI Titan Krios GII |
| Voltage (kV) | 300 |
| Camera | Gatan K2-Summit |
| Energy Filter | Gatan Quantum-LS (GIF) |
| Pixel size (Å/pix) | 1.059 |
| Preset target global defocus range (μm) | 0.5 - 3.5 |
| **3D reconstruction** | |
| Number of movies | 8882 |
| Initially selected particle candidates | 2 997 079 |
| Final number of particles | 326 257 |
| Resolution $_{\text{FSC independent half-maps}}$ (Å) | 4.3 |
| Local resolution range (Å) | 4.1 – 7.6 |
| Sharpening B-factor (Å$^2$) | -140.0 |
| **Refinement** | |
| Number of atoms | 30 333 |
| Residues | 1966 |
| Ligands | / |
| $CC_{box}$, $CC_{mask}$, $CC_{volume}$ (Afonine et al., 2018) | 0.82, 0.72, 0.72 |
| CCs of individual chains (Afonine et al., 2018) | |
| Receptor dimer (chain IDs): A, B, C, D | 0.74, 0.76, 0.74, 0.74 |
| Head-bound insulins: E, F; G, H | 0.76, 0.68; 0.78, 0.71 |
| Stalk-bound insulins: I, J; K, L | 0.78, 0.79; 0.79, 0.78 |
| Resolution $_{\text{FSC map vs. model@0.143}}$ (Å) (Afonine et al., 2018) | 3.79 |
| RMSD | |
| Bond lengths (Å) | 0.005 |
| Bond angles (°) | 0.964 |
| Ramachandran favored (%) | 82.3 |
| Ramachandran gen. allowed (%) | 17.7 |
| Ramachandran disallowed (%) | 0.0 |
| MolProbity score | 1.95 |
| Clash score | 4.8 |

**Table S2: Overview of residues included and absent from the IR-ECD model refined against the electron microscopic data.** Modeled residue ranges, domain assignments and residues with map-to-model correlation coefficients below 0.6 (as a measure of confidence) are listed. Furin cleavage recognition sites have been omitted from this model but included in the MD simulation model.

| Polypeptide name, chain in pdb (residue range) | Domain (residue range) | Residues with map-to-model cc ≤ 0.6 |
|---|---|---|
| IRα chain, chain A (1-719) | L1 (1-157) | Leu 2, Glu 24, Asn 25, Asn 111, Ile 112 |
| | CR (158-310) | Cys 182, Thr 184, Cys 201, Thr 202, Cys 207, Cys 208, Cys 212, Asn 215, Cys 216, Cys 266, Met 294 |
| | L2 (311-470) | Asn 397 |
| | FnIII-1 (471-595) | - |
| | FnIII-2α (596-637) | Gln 610 |
| | IDα (638-688) | Asp 645, Tyr 646, Lys 649, Phe 662, Glu 663, Ser 664, Ser 667, Gln 668, Lys 669, His 670, Glu 676, Asp 677, Ser 678, Ala 679, Gly 680 |
| | α CT (689-723) | Phe 714 |
| IRβ chain, chain B (756-917) | IDβ (724-755) | Absent from model |
| | FnIII-2β (756-808) | His 775, Phe 776, |
| | FnIII-3 (809-917) | Gly 840, Leu 844, Tyr 845, Glu 846, Val 847, Gly 853, Asp 854, Glu 855, Cys 860, Val 861, Ser 862, Gly 880, Asn 881, Asn 881, Tyr 882, Val 884, Ile 886, Arg 887, Ala 888, Thr 898, Phe 903, Val 905, Ser 913 |
| IRα' chain, chain C (1-719),<br><br>partial density for furin cleavage site (720-723) present in reconstruction | L1' (1-157) | Asn 15, Asn 25, Leu 37, Asn 111, Tyr 127, Asp 138 |
| | CR' (158-310) | Thr 172, Val 191, Cys 192, Cys 201, Thr 202, Cys 208, Cys 212, Asn 215, Cys 216, Arg 229, Cys 237 |
| | L2' (311-470) | - |
| | FnIII-1' (471-595) | Ser 573 |
| | FnIII-2α' (596-637) | Asn 606, Ser 607, Gln 610 |
| | IDα' (638-688) | Cys 647, Gly 650, Leu 651, Lys 652, Thr 657, Trp 658, Ser 659, Pro 660, Pro 661, Phe 662, Glu 663, Ser 664, Glu 665, Asp 666, Ser 667, Gln 668, Lys 669, His 670, Asn 671, Gln 672, Ser 673, Glu 674, Glu 676, Asp 677, Ala 679, Cys 683 |
| | αCT' (689-723) | - |
| IRβ' chain, chain D (756-917) | IDβ' (724-755) | Absent from model |
| | FnIII-2β' (756-808) | His 756, Phe 776, Arg 804 |
| | FnIII-3' (809-917) | Asn 826, Asn 839, Gly 840, Leu 841, Glu 846, Gly 853, Asp 854, Glu 855, Gly 871, Arg 875, Val 884, Arg 885, Ile 886, Gly 895, Val 905 |
| Insulin 1, chain E (1-21) | A-chain | Tyr 19 |
| chain F (1-30) | B-chain | Gly 23, Phe 25 |
| Insulin 1', chain G (1-21) | A-chain | - |
| chain H (1-30) | B-chain | Arg 22, Phe 25 |
| Insulin 2, chain I (1-21) | A-chain | - |
| chain J (1-30) | B-chain | - |
| Insulin 2', chain K (1-21) | A-chain | - |
| chain L (1-30) | B-chain | - |

**Table S3: Glycan composition of the IR-ECD MD simulation model.** N- and O-linked glycans are based on the analysis by Sparrow et al., 2007, and Sparrow et al., 2008.

| Residue (Monomer 1) | Residue (Monomer 2) | Glycan composition |
|---|---|---|
| **N-linked glycosylation sites** | | |
| Asn16 | Asn16’ | $GlcNAC_2Man_5$ |
| Asn25 | Asn25’ | $GlcNAC_4Man_3Gal_2Fuc$ |
| Asn111 | Asn111’ | $GlcNAC_2Man_9$ |
| Asn215 | Asn215’ | $GlcNAC_2Man_6$ |
| Asn255 | Asn255’ | $GlcNAC_4Man_3Gal_2Fuc$ |
| Asn295 | Asn295’ | $GlcNAC_4Man_3Gal_2Fuc$ |
| Asn337 | Asn337’ | $GlcNAC_2Man_5$ |
| Asn397 | Asn397’ | $GlcNAC_2Man_5$ |
| Asn418 | Asn418’ | $GlcNAC_4Man_3Gal_2Fuc$ |
| Asn514 | Asn514’ | $GlcNAC_2Man_5$ |
| Asn606 | Asn606’ | $GlcNAC_4Man_3Gal_2Fuc$ |
| Asn624 | Asn624’ | $GlcNAC_4Man_3Gal_2Fuc$ |
| Asn671 | Asn671’ | $GlcNAC_6Man_3Gal_4Fuc$ |
| Asn730 | Asn730’ | $GlcNAC_4Man_3Gal_2Fuc$ |
| Asn743 | Asn743’ | $GlcNAC_4Man_3Gal_2Fuc$ |
| Asn881 | Asn881’ | $GlcNAC_4Man_3Gal_2Fuc$ |
| Asn894 | Asn894’ | $GlcNAC_2Man_6$ |
| **O-linked glycosylation sites** | | |
| Thr732 | Thr732’ | $(GalNAC.Gal)_2$ |
| Thr737 | Thr737’ | $(GalNAC.Gal)_2$ |
| Ser745 | Ser745’ | $(GalNAC.Gal)_3$ |
| Ser746 | Ser746’ | $(GalNAC.Gal)_3$ |
| Thr747 | Thr747’ | $(GalNAC.Gal)_3$ |
| Thr751 | Thr751’ | $(GalNAC.Gal)_3$ |

**Table S4: Center-of-mass distances measured between insulin B-chain C-terminal and B-chain α-helix residues for 10 MD simulations (R1-R10).** The respective residues are indicated in Fig. S14. C-α distances between residues have been averaged for each simulation run over 500 ns simulation time.

| Insulin 1 | C-α distance in cryo-EM structure (Å) | C-α distance in MD simulations (Å) | | | | | | | | | |
|---|---|---|---|---|---|---|---|---|---|---|---|
| | | R1 | R2 | R3 | R4 | R5 | R6 | R7 | R8 | R9 | R10 |
| Phe B24 – Leu B15 | 8.7 | 10.24 | 10.74 | 13.32 | 12.61 | 8.40 | 9.71 | 10.38 | 9.41 | 6.96 | 11.51 |
| Phe B25 – Leu B15 | 11.49 | 13.32 | 14.25 | 15.52 | 15.32 | 11.92 | 13.25 | 13.32 | 12.40 | 9.41 | 14.33 |
| Tyr B26 – Val B12 | 13.73 | 16.43 | 16.82 | 16.79 | 18.02 | 13.84 | 16.64 | 15.49 | 14.79 | 13.39 | 16.55 |
| Thr B27 – Val B12 | 17.54 | 19.04 | 20.29 | 19.43 | 20.88 | 17.02 | 20.17 | 18.13 | 18.46 | 17.11 | 20.00 |
| Pro B28 – Gly B8 | 23.91 | 23.96 | 26.60 | 21.49 | 25.76 | 22.45 | 26.04 | 22.03 | 25.57 | 23.46 | 25.56 |
| Lys B29 – Gly B8 | 27.05 | 24.46 | 28.94 | 21.08 | 28.46 | 24.17 | 29.14 | 21.37 | 26.77 | 26.04 | 27.70 |
| Thr B30 – Gly B8 | 26.75 | 26.30 | 27.62 | 22.60 | 30.73 | 22.73 | 30.50 | 22.0 | 29.52 | 27.00 | 30.18 |

| Insulin 1' | C-α distance in cryo-EM structure (Å) | C-α distance in MD simulations (Å) | | | | | | | | | |
|---|---|---|---|---|---|---|---|---|---|---|---|
| | | R1 | R2 | R3 | R4 | R5 | R6 | R7 | R8 | R9 | R10 |
| Phe B24 – Leu B15 | 9.63 | 10.27 | 12.89 | 11.86 | 13.88 | 14.34 | 17.30 | 11.55 | 12.25 | 8.91 | 10.31 |
| Phe B25 – Leu B15 | 13.35 | 13.77 | 15.39 | 15.37 | 17.28 | 17.21 | 20.33 | 15.18 | 15.44 | 12.54 | 13.07 |
| Tyr B26 – Val B12 | 17.82 | 16.45 | 19.78 | 18.12 | 21.83 | 20.35 | 22.31 | 17.05 | 17.76 | 15.06 | 15.52 |
| Thr B27 – Val B12 | 21.23 | 19.01 | 22.51 | 20.36 | 24.50 | 22.31 | 25.84 | 20.39 | 18.95 | 18.72 | 17.56 |
| Pro B28 – Gly B8 | 27.47 | 26.73 | 30.85 | 24.85 | 33.04 | 32.15 | 32.81 | 27.40 | 21.85 | 23.53 | 22.21 |
| Lys B29 – Gly B8 | 27.12 | 27.25 | 33.65 | 26.25 | 35.32 | 34.30 | 33.68 | 27.67 | 24.72 | 24.59 | 22.82 |
| Thr B30 – Gly B8 | 28.12 | 29.27 | 35.36 | 27.00 | 37.46 | 37.15 | 35.04 | 30.01 | 25.81 | 27.42 | 25.00 |

| Insulin 2 | C-α distance in cryo-EM structure (Å) | C-α distance in MD simulations (Å) | | | | | | | | | |
|---|---|---|---|---|---|---|---|---|---|---|---|
| | | R1 | R2 | R3 | R4 | R5 | R6 | R7 | R8 | R9 | R10 |
| Phe B24 – Leu B15 | 6.5 | 9.83 | 7.27 | 8.32 | 8.22 | 8.07 | 8.32 | 8.00 | 6.79 | 7.12 | 6.97 |
| Phe B25 – Leu B15 | 7.77 | 11.43 | 9.12 | 11.30 | 10.96 | 10.30 | 8.79 | 11.14 | 8.32 | 10.04 | 8.97 |
| Tyr B26 – Val B12 | 7.15 | 10.10 | 6.37 | 11.64 | 11.77 | 9.82 | 7.32 | 13.43 | 8.19 | 8.81 | 7.20 |
| Thr B27 – Val B12 | 7.68 | 11.88 | 6.88 | 13.69 | 14.56 | 12.29 | 7.91 | 14.90 | 7.73 | 7.38 | 10.30 |
| Pro B28 – Gly B8 | 8.33 | 6.12 | 4.57 | 12.66 | 17.66 | 14.60 | 6.00 | 17.90 | 12.41 | 6.36 | 13.46 |
| Lys B29 – Gly B8 | 9.79 | 9.34 | 5.31 | 14.69 | 18.59 | 15.69 | 4.86 | 17.98 | 11.88 | 6.25 | 17.01 |
| Thr B30 – Gly B8 | 12.95 | 10.97 | 8.92 | 15.13 | 20.84 | 16.70 | 5.50 | 20.10 | 11.52 | 8.29 | 18.28 |

| Insulin 2’ | C-α distance in cryo-EM structure (Å) | C-α distance in MD simulations (Å) | | | | | | | | | |
|---|---|---|---|---|---|---|---|---|---|---|---|
| | | R1 | R2 | R3 | R4 | R5 | R6 | R7 | R8 | R9 | R10 |
| Phe B24 – Leu B15 | 7.63 | 8.45 | 8.55 | 8.18 | 8.77 | 8.04 | 7.98 | 8.00 | 8.27 | 9.45 | 7.80 |
| Phe B25 – Leu B15 | 9.26 | 10.95 | 10.90 | 10.68 | 11.30 | 10.01 | 10.31 | 10.69 | 11.23 | 12.11 | 9.89 |
| Tyr B26 – Val B12 | 7.03 | 12.75 | 9.16 | 9.42 | 8.94 | 9.09 | 8.23 | 9.24 | 11.23 | 12.90 | 7.76 |
| Thr B27 – Val B12 | 9.91 | 15.01 | 10.29 | 7.81 | 7.35 | 10.64 | 10.24 | 9.32 | 11.43 | 14.37 | 8.33 |
| Pro B28 – Gly B8 | 10.14 | 21.28 | 11.49 | 9.18 | 8.74 | 10.35 | 11.08 | 7.85 | 8.85 | 16.00 | 8.29 |
| Lys B29 – Gly B8 | 9.25 | 21.98 | 10.90 | 7.37 | 7.35 | 10.28 | 10.44 | 5.84 | 10.30 | 17.20 | 8.28 |
| Thr B30 – Gly B8 | 9.30 | 23.27 | 11.50 | 9.29 | 8.21 | 9.38 | 10.14 | 6.58 | 10.32 | 19.34 | 10.16 |
